# Development and Validation of Genome-Informed and Multigene-based qPCR and LAMP Assays for Accurate Detection of *Dickeya solani*: A Critical Quarantine Pathogen Threatening Potato Industry

**DOI:** 10.1101/2024.03.21.586178

**Authors:** Shefali Dobhal, Gem Santillana, Michael J. Stulberg, Dario Arizala, Anne M. Alvarez, Mohammad Arif

## Abstract

*Dickeya solani*, one of the most aggressive pectinolytic phytopathogens, causes blackleg disease in potatoes, resulting in significant economic losses and adversely impacting one of the world’s most important food crops. The diagnostics methods are critical in monitoring the latent infection for international trade of potato seed-tubers and in implementation of control strategies. Our study employed a whole-genome comparative approach, identifying unique target gene loci (LysR and TetR family of transcriptional regulators gene regions), design loop-mediated isothermal amplification (LAMP) and a multi-gene-based multiplex TaqMan qPCR assays for specific detection and differentiation of *D. solani*. Both methods underwent meticulous validation with extensive inclusivity and exclusivity panels, exhibiting 100% accuracy and no false positives or negatives. The LAMP method demonstrated the detection limit of 100 fg and 1 CFU per reaction using pure genomic DNA and crude bacterial cell lysate, respectively. The qPCR detection limit was 1 pg, 100 fg and 10 fg with Quadplex, triplex and singleplex, respectively. None of the assay showed any inhibitory effect after adding host DNA (in qPCR) or crude lysate (in LAMP). The assays proved robust and reproducible in detecting the target pathogen in infected samples, with the LAMP assay being field-deployable due to its simplicity and rapid results acquisition within approximately 8.92 minutes. The reproducibility was confirmed by performing the assay in two independent laboratories. These developed assays offer a robust, rapid, and reliable solution for routine testing, with applications in phytosanitary inspection and epidemiological studies.

**IMPORTANCE:** *Dickeya solani*, one of the most aggressive soft rots causing-bacteria and a quarantine pathogen, poses a severe threat to food security by causing substantial economic losses to potato industry. Accurate and timely detection of this bacteria is vital for monitoring latent infections, particularly for international trade of potato seed tubers, and for implementing effective control strategies. In this research we have developed a LAMP and a multi-gene-based multiplex TaqMan qPCR assays for specific detection of *D. solani*. These assays, characterized by their precision, rapidity, and robustness, are crucial for distinguishing *D. solani* from related species. Offering unparalleled sensitivity and specificity, these assays are indispensable for phytosanitary inspection and epidemiological monitoring, providing a powerful tool for management of this threatening pathogen.

## Introduction

Members of the *Dickeya* genus, which are Gram negative necrotrophic Soft Rot *Pectobacteriaceae* (SRP) phytopathogens, are primarily responsible for the increased economic losses in the agricultural crops and ornamentals plants globally (Kado et al., 2006; Adeolu et al., 2016; Lisicka et al., 2018). Global losses of up to 30% (Agrios, 2007), estimated at US$50–100 million every year, have been reported for vegetables, fruits and ornamental plants (Pérombelon and Kelman, 1980; Pérombelon, 2002; Ma et al., 2007). The ability of these plant pathogenic species to infect a wide range of plant host species with wide geographic distribution poses a threat to agricultural biosecurity and is of great concern for global food security. These SRP members are causative agents of blackleg symptoms on potatoes and soft rot diseases in many other host plant species, including vegetables, fruits, and ornamentals. The virulence is mainly attributed to the production and secretion of an array of plant cell wall-degrading enzymes, resulting in breakdown of plant tissues and release of nutrients that support bacterial growth (Arif et al., 2022; Boluk et al., 2021; Arizala and Arif, 2019; Alic et al., 2019; Charkowski et al., 2012; Joko et al., 2014; Reverchon et al., 2013). The genus *Dickeya* was originally established by Samson et al. (2005) with description of *D. chrysanthemi*, *D. paradisiaca, D. dadantii*, *D. dieffenbachiae, D. dianthicola*, and *D. zeae*. Brady et al. (2012) reclassified members of *D. dieffenbachiae* as a subspecies of *D. dadantii*. To date, this genus comprises 12 recognized and validly published names (https://www.bacterio.net/genus/dickeya; Arif et al., 2022), with the most recently described species being *D. lacustris, D. undicola, D. oryzae*, and *D. poaceiphila* (Hugouvieux-Cotte-Pattat *et al*., 2019; Oulghazi et al., 2019; Wang et al., 2020; Hugouvieux-Cotte-Pattat et al., 2020; Arif et al, 2022). The recent reclassification of the species placed in *D. paradisiaca* has proposed a reassignment to the genus level and named it *Musicola paradisiaca* (Hugouvieux-Cotte-Pattat et al., 2021). Among the species associated with this genus, *Dickeya solani*, first isolated in Poland in 2005, has emerged and is labeled as one of the most aggressive soft rot pathogenic bacteria. In potato, *D. solani* is responsible for blackleg disease in the fields and soft rot of tubers during the storage and transit. *Dickeya solani* can cause infection with lower inoculum, able to withstand a wide range of temperature from low to high, up to 39°C, which is favorable for disease development, out-competing other members of this genus. *Dickeya solani* has been reported to be present in Europe, Israel Brazil, and Turkey (Tsror et al., 2009; Toth et al., 2011; Van der Wolf et al., 2014; Cardoza et al., 2016; Ozturk and Aksoy, 2017). It has not yet been reported or known to be established in the United States. The dissemination of the pathogen across borders has been reported through the trade of infected vegetative propagating material and latently infected seed tubers, resulting in serious economic losses (Tsror et al., 2009; Toth et al., 2011). The aggressiveness of *D. solani* is reported to be enhanced with ambient temperature and increased rainfall, resulting in flooding in the fields, allowing the spread between the plants (Toth et al., 2011). The pathogen invades the vascular tissue, and spread has been reported from mother to daughter tubers (Czajkowski et al., 2010). *Dickeya solani* has been reported as a clonal population (Laurila et al., 2008; Slawiak et al., 2009; Van Vaerenbergh et al., 2012; Tsror (Lahkim) et al., 2009; van der wolf et al., 2014). Recently, whole-genome comparative analysis of *D. solani* has also been conducted, revealing a high level of homogeneity (Motyka-Pomagruk et al., 2020). To date, no resistant varieties or efficient chemicals have been reported to control the disease. Only crop rotation, careful handling during cutting, loading, and harvest, and removal of the diseased plants are recommended. Accurate and early-stage detection of the pathogens facilitates effective intervention and prevents further dissemination of the contaminated tubers and stolons.

PCR-based techniques are more sensitive, specific, and possess discriminatory abilities compared to immunological assays. Furthermore, real-time qPCR offers greater speed, sensitivity, and real-time amplification compared to endpoint PCR (Kim et al., 2008; Arif et al., 2013, 2014). TaqMan-probe-based qPCR amplification formats offer more specific and sensitive amplification. The use of a multi-gene approach to target two unique regions in the genome provides increased reliability, specificity, and minimizes the risk of false positives or false negatives (Arif et al., 2013).

However, the high costs and sophisticated laboratory equipment requirements often limit the use of real-time qPCR methods for field surveys and routine pathogen screenings. Currently, isothermal-based methods are becoming more popular due to their adequacy for point-of-care applications, sensitivity, fast results interpretation, and minimal equipment requirements. Loop-Mediated Isothermal Amplification (LAMP), originally reported by Notomi et al. (2000), is an auto cycling and strand displacement amplification method operating at a constant temperature (60–65°C), which can be accomplished using a water bath and obviates the need of a sophisticated thermocycler. It utilizes four to six specially designed LAMP primers targeting six to eight regions of the template DNA, ensuring high specificity (Nagamine et al., 2002). High sensitivity, specificity, rapidity, and comparatively high tolerance to matrix inhibitors make it an indispensable tool for on-field amplification method (Kaneko et al., 2007; Notomi et al., 2000; Arif et al, 2021). Melt curve analysis (MCA) is an approach for assessing the dissociation properties of nucleic acid amplicons based on their thermal stability. MCA, in conjunction with the LAMP method, is a rapid method for differentiating pathogenic agents, contamination, and non-specific products (Tone et al., 2017). LAMP can be easily deployed in non-specialized laboratories for conducting routine diagnostics through visual observation (Dobhal et al., 2020).

Current methods for the specific detection of *D. solani* include the TaqMan qPCR developed by Pritchard et al. (2013), the method based on the *fliC* gene by Van Vaerenbergh (2012), and the approach by Kelly et al. (2012) based on the *fusA* gene. Ivanov et al. (2020) reported a recombinase polymerase amplification (RPA) method, which utilizes primers modified or adapted from the previously reported SOL-C primers by Pritchard et al. (2013).

Currently, there are no LAMP or multi-gene based TaqMan real-time qPCR assays reported for *D. solani* that can be used in routine diagnostics or epidemiological surveys for quarantine applications. Furthermore, there is a pressing need for rapid and validated field-deployable diagnostic tools, especially to intercept the invasion of exotic pathogens. In this paper, we describe the development and validation of a genome informed, sensitive, fast, and reliable LAMP, as well as a multi-gene-based Real-time PCR (qPCR) assays for accurate detection and differentiation of *D. solani*. To enhance the reliability of the qPCR method, we multiplexed (4-plex) *D. solani* primers with our previously developed *Dickeya* genus primers (Dobhal et al., 2020) and the universal internal control (UIC; Ramachandran et al., 2021). For diagnostic specificity, we demonstrated the application of this method by testing all known species of *Dickeya*. The developed methods will enhance the analytical capabilities of plant disease diagnostic laboratories, phytosanitary agencies, surveillance, and management agencies, thereby aiding in the prevention of the introduction and dissemination of soft rot bacteria.

## Materials and Methods

### Bacterial strains and DNA extraction

The targeted bacterial pathogen, *D. solani*, and phylogenetically related bacterial species, as well as the outgroup species and common host colonizers found in the close niche of the pathogen were used to develop and validate the specificity of the assays. The strains were selected to represent different genetic and geographic diversity (countries) comprising of a total 133 and 123 strains for validation of LAMP and multiplex TaqMan qPCR assays, respectively, are listed in Supplement Table 1. Bacterial strains prefixed with “A” and “PL” were obtained from Pacific Bacterial Collection and Phytobacteriology lab collection (University of Hawaii at Manoa) (stored at − 80 °C). The other bacterial type cultures and recently described *Dickeya* and *Pectobacterium* strains were procured from international culture collections, including BCCM/LMG (Belgian Co-ordinated collections of Micro-organisms, Belgium), CIRM-CFBP (CIRM-Plant Associated Bacteria, France) and ICMP (International Collection of Microorganisms from Plants, New Zealand). The strains were cultured by streaking on TZC media (2,3,5-triphenyl-tetrazolium chloride medium: peptone 10 g/l, dextrose 5 g/l, 0.001% TZC and agar 17 g/l) (Norman and Alvarez, 1989). The plates were incubated at 26°C (±2°C) overnight; single colonies were streaked again on TZC medium and incubated at 26°C (±2°C) overnight. The lyophilized cultures from LMG, ICMP and CFBP were revived on recommended media provided by the culture collections instructions. The DNA was isolated using DNeasy Blood and Tissue Kit (Qiagen, Germantown, MA) according to manufacturer’s instructions. The DNA concentrations were estimated using NanoDrop 2000c UV-Vis Spectrophotometer (Thermo Fisher Scientific Inc., Worchester, MA). For sensitivity assays, the genomic DNA was quantified using Qubit 4 spectrophotometer (Thermo Fisher Scientific, Waltham, MA). The strain’s identity was confirmed by sequencing the chromosomal replication initiation protein *dnaA* gene region of different bacterial species either in this study or in previous studies (Ahmed et al., 2018; Dobhal et al., 2019; Dobhal et al., 2020) (Supplement Table 1)

### Plant Growth, Inoculation, and Total Plant DNA Extraction

The plant samples used for validation were naturally infected plant samples from the field, artificially inoculated plants grown in the greenhouse as well as artificially inoculated tuber slices. Briefly, plants grown in controlled greenhouse for inoculations, four-week-old healthy potato plants were stab inoculated with different *Dickeya* and *Pectobacterium* species except for *D. solani,* which were inoculated onto potato slices. For plant inoculations, stem was stab inoculated 10 cm from above the ground with 100 µl of 10^8^ CFU/ml of bacteria or water (control) at an angle of 45° using a 200 µl of pipette tip, secured by wrapping a parafilm to prevent leakage to soil (Czajkowski et al., 2010). Inoculated plants were covered individually with plastic bags to maintain the humidity, tied and incubated at 28 to 30°C for 24h. Blackleg symptoms were observed at the base of the stem after 3 days post inoculation. Healthy plants inoculated with sterile water showed no symptoms. The DNA was extracted from the healthy and greenhouse inoculated plants using DNeasy Plant Mini Kit (Qiagen), according to manufacturer’s instructions with an additional step of using a Mini-Bead Beater 16 Center Bolt (Biospec products, Bartlesville, OK) for one minute at maximum speed to thoroughly rupture the plant cells. Similarly, DNA was isolated from asymptomatic and symptomatic field samples using the same protocol described above. For TaqMan qPCR and LAMP validation and optimization: 1 µl of isolated DNA was used. For on-site detection using LAMP, OptiGene Plant Material Lysis kit (OptiGene Limited, Horsham, UK) was used to extract crude DNA following the manufacturer’s instructions (Domingo et al., 2021). Briefly, about 80 mg of plant tissue (stem)/potato tuber was placed in a 5 ml tube with an iron ball and 1 ml lysis buffer. The tube was shaken vigorously manually for 1 min to ground the plant material. This plant lysate was transferred using a loop (10 μl) into a vial containing 2 ml dilution buffer and mixed; 5 μl from this dilution buffer was used as template in LAMP assay.

### Genome comparison, Target gene selection and primer design for LAMP and TaqMan-qPCR

Genomes of *Dickeya solani* strain D s0432-1 (NZ_CP017453.1), strain IPO2222 (NZ_CP015137.1), strain PPO9019 (NZ_CP017454.1), strain RNS 08.23.3.1.A (NZ_CP016928.1), IFB 0099 (NZ_CP024711.1); *D. chrysanthemi* Ech1591 (NC_012912), *D.dadantii* strain 3937 (NC_014500), *D. dianthicola* strain RNS04.9 (NZ_CP017638.1), *D. fangzhongdai* DSM101947 (NZ_CP025003), *D. zeae* strain Ech586 (NC_013592), *Musicola paradisiaca* NCPPB2511 previously called as *D. paradisiaca* strain, *D. aquatica* 174/2 (NZ_LT615367.1), *Pectobacterium carotovorum* strain PCC21 (NZ_018525), *Erwinia amylovora* strain CFBP1430 (NC_013961), *Ralstonia solanacearum* GMI 1000 (NC_003295), and *Clavibacter sepedonicus* strain ATCC33113 (NC_10407) were retrieved from NCBI GenBank Genome database. Whole genomes were aligned using progressive Mauve (2.4.0) and Geneious (10.2.3) (Darling et al., 2020). Generated Locally Collinear Blocks (LCBs) were examined to identify unique and conserved regions within *D. solani.* The unique *tetR* family transcriptional regulator gene specific for *D. solani* was used for designing primers for LAMP (Table 1). For multiplex qPCR detection of *D. solani,* we employed both *tetR* gene and an additional distinct target gene region from the *lysR* family transcriptional regulator gene to design primers and probes (Table 2). The representative genome of each *Dickeya* species and complete genomes of *D. solani* along with the genomes of other closely related genera and selected target gene locations were included to generate a BLAST Ring Image Generator (BRIG) 40 (Alikhan et al., 2011). NCBI-BLAST 2.6.0+ database was used to compare and generate the BRIG image.

**Table 1.**
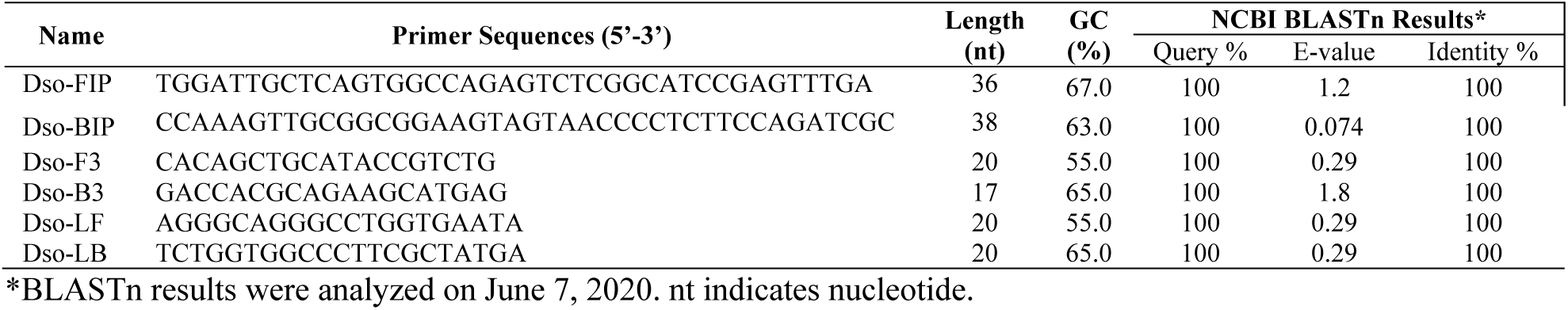
Primers developed for loop-mediated isothermal amplification (LAMP) assay for specific detection of *Dickeya solani* targeting tetR family transcriptional regulator gene.

**Table 2.**
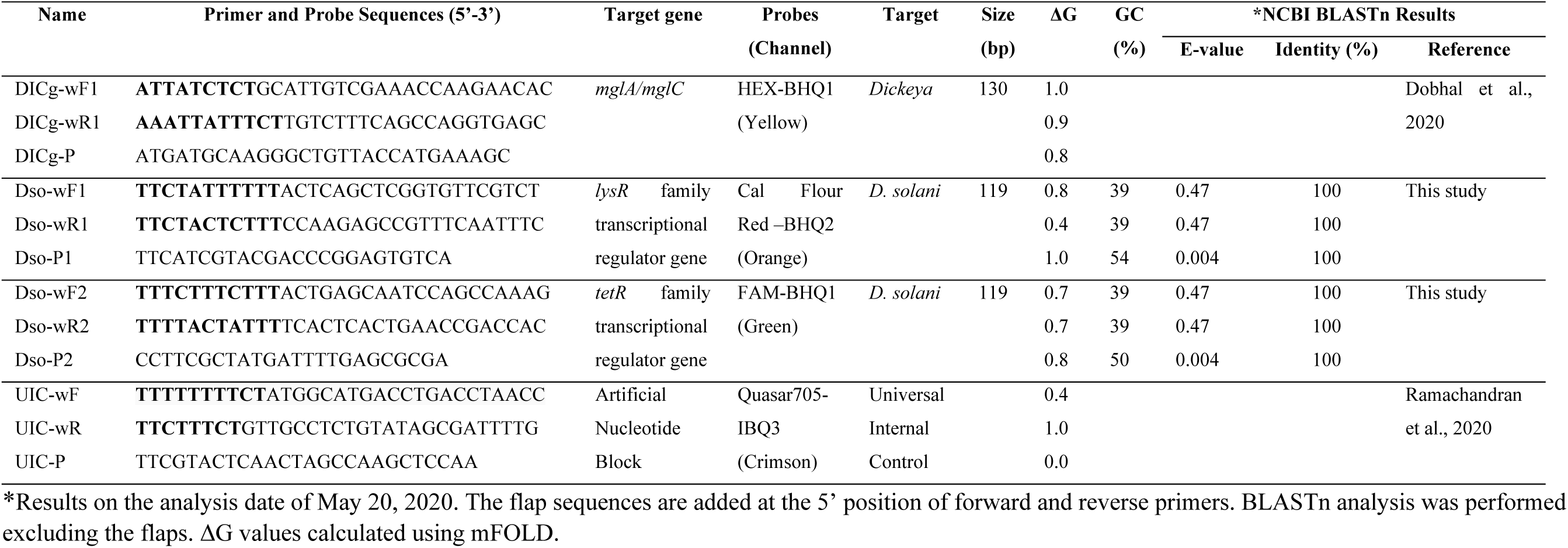
Primers and probes information for 4-plex qPCR assay developed for specific detection of *Dickeya solani*.

The unique loci for *tetR* family transcriptional regulator gene specific for *D. solani* was used to develop LAMP primers that amplify all isolates within this species. The LAMP primers, forward inner primer (FIP), backward inner primer (BIP), outer primers (F3 and B3) were designed using PrimerExplorer V5 software (Eiken Chemical Company; https://primerexplorer.jp/e/). Internal loop primers, forward loop primer (LF) and backward loop primer (LB) were designed manually to accelerate the LAMP reaction. The primers were synthesized by Genewiz from Azenta Life Sciences Inc (Genewiz, Inc., South Plainfield, NJ). The specificity of each primer was confirmed by comparing the primer sequences using the NCBI GenBank BLASTn tool. Primers were checked for their thermodynamic parameters as described by Arif and Ochoa-Corona (2013).

For multiplex TaqMan qPCR, for enhanced reliability two sets of primers targeting two loci *lysR* and *tetR* family transcriptional regulator gene, unique regions for *D. solani* species, were selected and used for primer and probe design with Primer3 software (Rozen and Skaletsky, 2000). The primers and probes were evaluated *in silico* for thermodynamic characteristics, internal structures and self-dimer formations with mFold (Zuker, 2003; Arif and Ochoa-Corona, 2013). The specificity of the primers and probes were confirmed *in silico* by screening the primers and probes sequences using BLASTn tool with the NCBI GenBank nucleotide and genome databases. Furthermore, to adjust Tm, GC content and the length of the primers, customized flap sequences were added to the 5’ end of each primer as described previously (Arif and Ochoa-Corona, 2013; Larrea et al., 2019). For multiplexing, the combinations of fluorophore and quencher were used to detect two different loci of the pathogen *D. solani*: Cal Flour Red –BHQ2 (*LysR* target gene 1), FAM-BHQ1(*tetR* target gene 2) which were multiplexed with previously described HEX-BHQ1 (*Dickeya* genus, Dobhal et al., 2020) and Quasar705-IBQ3 (universal internal control, Ramachandran et al., 2020). The primers and probes sequences along with the thermodynamic parameters are presented in Table 2. All primers and quencher probes 5’-/6FAM/BHQ-1/-3’ and 5’-/CAL Fluor Red/BHQ-2/-3’ were synthesized by Genewiz Inc. and LGC, Biosearch Technologies (LGC, Biosearch Technologies, Petaluma, CA). The universal internal control primers and probes were included in the reaction to serve as reagent control and to evaluate the presence of any PCR inhibitors in the reaction. The annealing temperatures, extension temperatures of primers and probes as well as PCR specificity were determined by first testing each individual primers/probe set with an endpoint PCR, followed by qPCR assay, and then confirming the parameters in the multiplex qPCR assay. All oligonucleotide sequences used for the LAMP and multiplex TaqMan-qPCR are listed in Tables 1 and 2.

### LAMP: Assay validation specificity and sensitivity

LAMP reactions (final volume of 25 µl) were performed in Rotor-Gene Q (Qiagen, Germatown, MD). The reaction mixture contained 15 µl of Optigene Master Mix (Optigene, West Sussex, United Kingdom), 2 µl primer mix containing 1.6 μM of each Dso-FIP/BIP, 0.2 μM of each Dso-F3/B3, 0.4 μM of each Dso-LF/LB, 1 µl of template DNA and 7 µl of nuclease free water (Invitrogen). The reaction mixture was incubated and amplified using Rotor-Gene Q at 65°C for 20 min followed by melt curve analysis at 99-80°C with an increment of 0.2 °C/ s. A reaction with no template DNA (nuclease free water) and DNA from a healthy plant were used as negative controls; *D. solani* DNA was used as a positive control. Positive and negative controls were included in all assays. The fluorescence was monitored in real-time using Rotor-Gene producing sigmoid shape curve on the generated plot, and amplified product was also verified by adding 3 µl of SYBR Green I dye (1:9 dilution, Life technologies Corporation, Eugene, OR) in each sample tube. The color of positive amplification products turns fluorescent green while those remaining orange were considered negative. Results with SYBR Green dye were visualized directly either with the naked eyes and or under ultraviolet light and using in gel documentation system (Bio-Rad Gel Doc-XR+, Bio-Rad). In addition, 10 µl of LAMP product was electrophoresed on 2% agarose gel with 1X TAE buffer, stained with ethidium bromide and visualized in gel documentation system. A ladder-like amplification was considered as a positive reaction for LAMP assay.

For assay specificity, DNA templates from 133 bacterial strains were tested (Supplement Table 1). The specificity assay comprised of inclusivity-11 strains of *D. solani* from eight different geographic locations worldwide; exclusivity panel consist of 122 strains of phylogenetically closely related plant pathogenic gram-positive and gram-negative bacterial species, shared niche bacterial species from different hosts and geographic locations, and endophytes isolated from the potato samples, and healthy plant DNA (Supplement Table 1).

The sensitivity of the LAMP assay was determined with both ten-fold serially diluted genomic DNA (A5581) and DNA from heat killed bacterial culture (serially diluted from 10^8^ to 1 CFU/ml). For sensitivity using the genomic DNA, ten-fold serially diluted pure genomic DNA, 10 ng to 1 fg, of *D. solani* were prepared in nuclease free water. For sensitivity with bacterial cells, overnight grown culture of *D. solani* (ca. 10^9^ CFU/ml) were ten-fold serially diluted to 1 CFU/ml in 0.1% sterile peptone water (BD, Becton Dickinson). The bacterial concentration was enumerated by spread plating in triplicate 100 *μ*l from serially diluted 10^-6^, 10^-7^ and 10^-8^ dilutions in 0.1% sterile peptone water on TZC media and plates were incubated at 28°C for 12-18 h. The microbial colonies were enumerated, and the average of microbial counts was reported as log_10_ CFU/ml. For performing the sensitivity, serially diluted bacterial cells were heat killed at 95 °C for 10 mins and centrifuged. One microliter of the serially diluted DNA samples from each dilution series (for both genomic and bacterial cells) was added into individual LAMP reaction mixture. Additionally, detection limit of the developed assay was also performed in presence of host genomic DNA for both spiked LAMP assays using genomic DNA and bacterial cells. OptiGene Plant Material lysis kit (OptiGene) was used to extract the host plant DNA as described earlier, and 5 μl from this was used for LAMP spiked assay. A non-template control (NTC; water) was included in all LAMP assays. These dilutions along with a non-template control were amplified in a Rotor-Gene Q Thermocycler; the LAMP conditions and method were followed as described above.

### Multiplex-TaqMan qPCR: Assay specificity and sensitivity

The multiplex real-time qPCR assays were carried out in Rotor-Gene Q (Qiagen). A primer mix (200 µl) was prepared by adding 10 µl of each forward and reverse primer (100 µM stock) targeting the genus *Dickeya* (Dobhal et al., 2020), *D. solani* (2 targets) and internal control (UIC) (Ramachandran et al., 2021) and 120 µl nuclease free water. The amplification was performed in a 25 μl reaction volume containing 12.5 μl of Rotor-Gene Multiplex PCR Master Mix (Qiagen), 2 μl of the primer mix (5 µM each primer concentration: DICg-wF1, DICg-wF1R1; Dso-wF1, Dso-wR1; Dso-wF2,wR2; UIC-wF, UIC-wR), 0.5 μl of each TaqMan probe (DICg-P, Dso-P1, Dso-P2 and UIC probes: 5 µM stock), 1 µl of plasmid DNA (Universal internal control, UIC-1 pg; Ramachandran et al., 2021), 1 µl of template DNA and nuclease free water was adjusted to obtain final volume. Positive and negative controls (non-template; water) were included in each TaqMan qPCR amplification run. The amplification conditions included an initial denaturation step at 95°C for 5 min, followed by 40 cycles, each one consisting of 95 °C for 30 s, and 60 °C for 15 s, acquiring fluorescence on both yellow (HEX), orange (Cal Flour Red), green (FAM), and crimson (Quasar705) channels at the end of each extension steps. Each qPCR reaction was performed in three replicates; standard deviation was calculated. The data analysis was done using the Rotor-Gene Q series software 2.3.1 (Built 49) with auto threshold (Ct); dynamic tube-based normalization was used. All threshold cycle (C_T_) values ≤35 was classified as a positive reaction. Additionally, the specificity was performed using the genomic DNA of 123 strains, including *D. solani* (n=11), closely and distantly related bacterial species and the endophytes isolated from the potato plants (n=112) similarly as indicated in LAMP assay. The multiplex TaqMan qPCR was performed using the same conditions as indicated above. Each strain was run in three replicates; standard deviation was calculated. The inclusion of universal internal control showed fluorescent signals which indicates any PCR inhibition in the reaction.

The detection limit (sensitivity) of the developed multiplex TaqMan-qPCR was evaluated by generating standard curves for the multiplex and singleplex qPCR. A 10-fold serial dilution from 10 ng to 1 fg of pure genomic DNA of *D. solani* (A5581) was prepared in nuclease free distilled water, and 1 μl of each dilution was used as template. For comparison, five qPCR sensitivity assays were performed: (1) Four-plex qPCR assay using all four primers/probes (DICg-wF1/DICg-wF1R1/DICg-P; Dso-wF1/Dso-wR1/Dso-P1; Dso-wF2/wR2/Dso-P2; UIC-wF/UIC-wR/UIC-P) together; (2) three-plex using 3 primer/probe sets (DICg-wF1/DICg-wF1R1/DICg-P; Dso-wF1/Dso-wR1/Dso-P1; Dso-wF2/wR2/Dso-P2) without the universal internal control; (3) single-qPCR using Dso-wF1/wR1/Dso-P1 primers only targeting *D. solani lysR* family transcriptional regulator gene only; (4) single-qPCR using Dso-wF2/wR2/Dso-P2 primers targeting *D. solani tetR* family transcriptional regulator gene only; and (5) spiked sensitivity assay with all four sets as indicated above plus 1 µl of healthy host (potato) genomic DNA. In spiked assay, host DNA was added in each qPCR reaction mix containing 10-fold serially diluted genomic DNA of *D. solani* (A5581) from 10 ng to 1 fg. The concentration of 1 pg was consistently used in the multiplex qPCR assays. Each reaction was performed in three replicates and the real-time qPCR assay was performed in the Rotor-Gene Q using the conditions described above. The genes used to develop these assays were present as a single copy in *D. solani* genome. Therefore, based on the genome size (estimated genome size of *D. solani is* ~ 4.84-4.96 Mb), the copy number was calculated by the web-based software (www.scienceprimer.com/copy-number-calculator-for-realtime-pcr or http://cels.uri.edu/gsc/resources/cndna.html).

### Validation with naturally, and artificially inoculated samples, and multi-lab-based validation

To assess the applicability of the developed assays, validation was conducted using naturally infected samples (collected from the field) and artificially inoculated samples. Field samples were collected from the Oahu fields in Hawaii, January 2019 (Arizala et al., 2020). Briefly, potato plants showed symptoms for blackleg were brought to the lab and processed, stem samples were cut 1 cm in size, surface sterilized 0.6% sodium hypochlorite for 30 s, followed by three rinses in sterilized distilled water. The DNA was extracted using DNeasy Blood and Tissue Kit (Qiagen). The plant samples infected with *Dickeya* and/or *Pectobacterium* were confirmed using previously developed assay in our lab (Dobhal et al. 2020, and Arizala et al 2022); the bacteria were isolated from these samples using CVP media (Helias et al., 2012), and incubated for 48 h at 26 ± 2°C. The colonies producing pits on CVP media, peculiar characteristics of pectinolytic bacteria, were picked and streaked to isolate single colonies and purified. The DNA was further isolated from the bacteria using DNeasy Blood and Tissue Kit (Qiagen), and the identity was further confirmed by sequencing *dnaA* (Dobhal et al. 2020). One microliter of DNA from infected plant collected from the fields (confirmed for the *Dickeya* and *Pectobacterium* species) were used for the validation of LAMP and multiplex TaqMan qPCR assays.

Further validation of both assays with known artificially inoculated samples, healthy “disease free” potato tubers were inoculated with different strains of *Dickeya* species. Briefly, the tubers were rinsed in water, surface sterilized with 0.6% sodium hypochlorite for 5 mins, rinsed with sterile water three times. The tuber slices were cut in the biosafety cabinet and placed into petridishes. Hundred microliters of 10^9^ CFU/ml of each species were inoculated on tuber slices, sterile water was inoculated for negative control. The plates were incubated at 28 °C ±2 °C in an incubator for 24 h. For multiplex TaqMan qPCR assay, the samples were processed using the DNeasy Plant Mini Kit (Qiagen), and 1 µl of DNA was used as a template. For LAMP assay, DNA was isolated using Plant Material Lysis Kits (Optigene) as per the manufacturer’s instructions. The amount of 5 μl crude DNA was added to LAMP reaction. Positive and negative controls were included in all assays.

The multi-lab validation test was performed in two different laboratories. Eight known bacterial genomic DNA were blind-coded and tested in both laboratories with the optimized LAMP and multiplex TaqMan qPCR protocols (as described above). The blind test panel comprised: *D. solani* (LMG25990, LMG27549, LMG27553, LMG27554), *D. zeae* (A5410), *D. dianthicola* (A5570), *P. parmentieri* (LMG29774), healthy potato (negative template control).

## Results

### Target selection, primers and probes design for TaqMan qPCR and LAMP assays

Highly conserved region within the core genome of *D. solani* was found by evaluating 16 genomes in the genus *Dickeya*, *Pectobacterium*, *Erwinia*, *Ralstonia*, *Clavibacter* and other closely related bacterial species using BLAST comparison and MAUVE (Figure 1). The BLAST comparison of whole genomes of *D. solani* computed using BRIG (Blast Ring Image Generator) (Alikhan et al., 2011), illustrated high degree of homogeneity between *D. solani* genomes. The TaqMan qPCR assay was designed using the LysR- and TetR family transcriptional regulators, which are common prokaryotic DNA binding proteins. These regulators consist of an N-terminal helix–turn–helix DNA-binding domain and a C-terminal co-inducer binding domain (Maddocks and Oyston, 2008; Orth et al., 2000; Sheehan et al., 2015). These regions were exclusively found in the genomes of *D. solani* and were absent in related species. LysR-type regulatory proteins are conserved across various bacterial species and control the expression of genes related to virulence, motility, quorum sensing, and metabolism (Doty et al., 1993; Heroven and Dersch, 2006; O’Grady et al., 2011; Hartmann et al., 2013, Sheehan et al., 2015). The TetR family transcriptional regulators primarily regulate genes involved in various catabolic pathways, antibiotic biosynthesis, multidrug resistance, osmotic stress response, and pathogenicity in both gram-positive and gram-negative bacteria (Ramos et al., 2005). For multiplex TaqMan qPCR, we targeted both target genes *tetR* and *lysR* gene. To improve assay sensitivity and primer compatibility for multiplexing, we incorporated 5’ tails, also known as flap sequences, into the primer sequences. Primers and probes designed for multiplex TaqMan qPCR were analyzed by BLAST analysis using NCBI nucleotide/genome database showed 100% query coverage and 100% identity only with the corresponding target sequences of *D. solani* (Table 2). The internal structures predictions and delta G (ΔG) plot values of each primer and probe were calculated using mFold (Table 2). All primers and probes had a ΔG of ≤1.0 at 60°C. The primers designed from the two distinct region of *D. solani*, incorporate flap sequences, enabling their multiplexing with previously designed genus *Dickeya* primers and universal internal control primers, enhancing the overall reliability of the assay.

**Figure 1:**
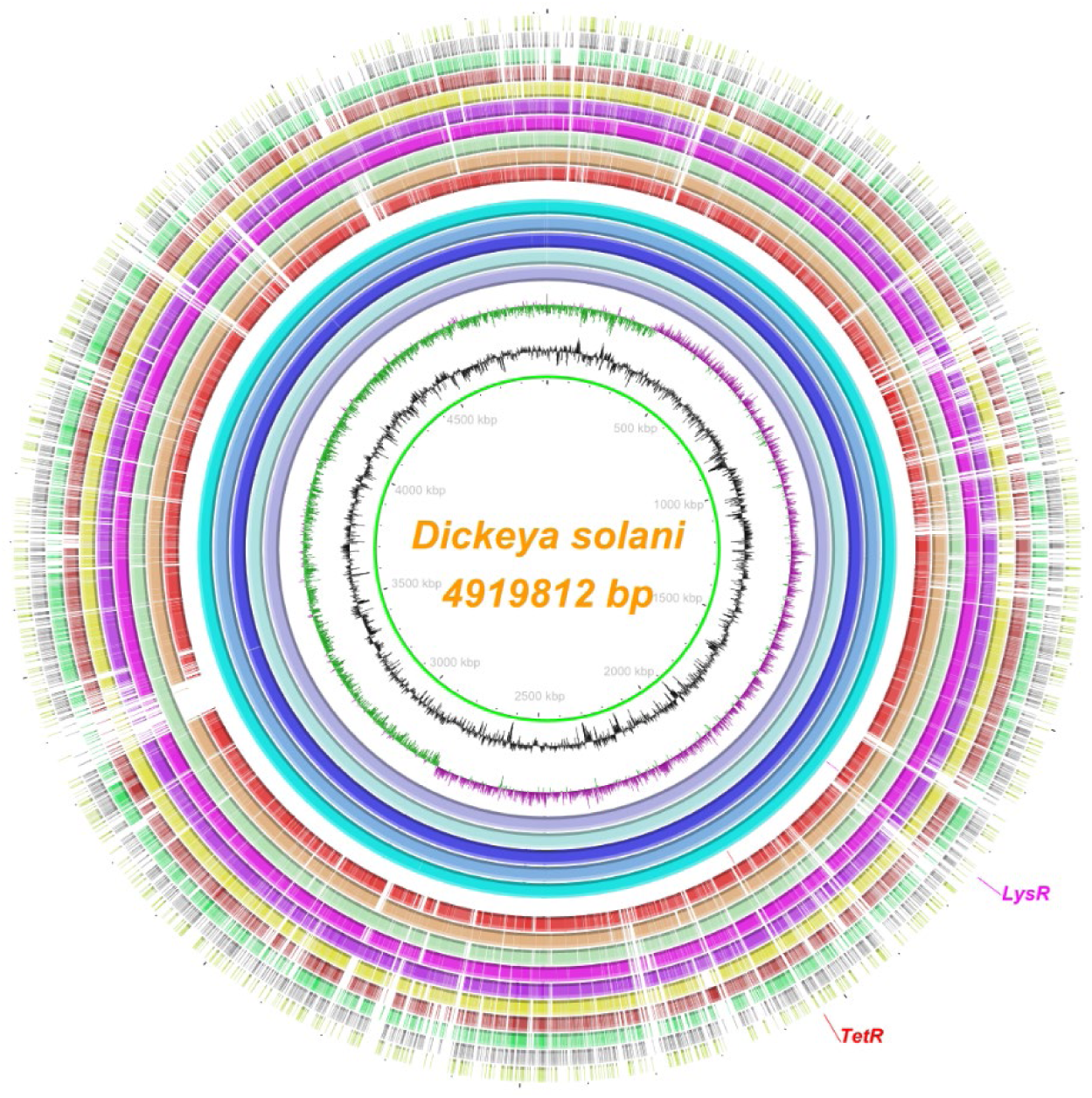
Ring plot representing the position of the two unique coding sequences TetR family transcriptional regulator and LysR family transcriptional regulator used for the specific detection of *Dickeya solani*. The layers show the multiple genome alignment of five *D. solani* strains followed by the other six members within the *Dickeya* genus and other bacterial species causing diseases to potato. From innermost to the outer, the layers indicate the genome coordinates (mega base pairs; mbp), the GC content (zigzag black line) and the GC skew (zigzag purple + / green −) of the reference genome *D. solani* type strain D s0432-1ᵀ. The color-coded rings illustrate from inwards the BLASTn pairwise comparison of *D. solani* D s0432-1ᵀ (NZ_CP017453.1), *D. solani* IPO2222 (NZ_CP015137.1), *D. solani* PPO9019 (NZ_CP017454.1), *D. solani* RNS 08.23.3.1.A (NZ_CP016928.1), *D. solani* IFB 0099 (NZ_CP024711.1), position of the two targets coding sequences LysR family transcriptional regulator and TetR family transcriptional regulator conserved across all *D. solani* strains (highlighted and labeled with fuchsia and red colors, respectively), *D. chrysanthemi* Ech1591ᵀ (NC_012912), *D. dadantii* 3937 (NC_014500), *D. parazeae* Ech586 (NC_013592; formerly known as *D. zeae*), *Musicola paradisiaca* Ech703 (NC_012880; previously called as *D. paradisiaca*), *D. dianthicola* RNS04.9ᵀ (NZ_CP017638.1), *D. fangzhongdai* DSM101947ᵀ (NZ_CP025003), *D. aquatica* 174/2ᵀ (NZ_LT615367), *P. carotovorum* PCC21 (NZ_018525), *Erwinia amylovora* CFBP 1430ᵀ (NC_013961), *Ralstonia pseudosolanacearum* GMI 1000ᵀ (NC_003295), and *Clavibacter sepedonicus* ATCC 33113ᵀ (NC_10407). The image was created using the BLAST Ring Image Generator (BRIG) v 0.95 (Alikhan et al., 2011).

Both target genes for *D. solani* demonstrated with 100% efficiency in the preliminary screening of the primers for their use in the LAMP assay; *tetR* family transcriptional regulator gene, showed lower Ct values when compared to the *lysR* gene, was used for designing the primers for LAMP assay. The *tetR* family transcriptional regulator gene analyzed by BLASTn using NCBI database showed 100% query coverage and 100% identity only with the corresponding target sequences of *D. solani*. *In-silico* analysis demonstrated no amplification with non-target genus *Dickeya*, *Pectobacterium*, *Clavibacter* and other closely related species, sharing the common niche spaces together. The length of primers and GC content varied from 17- to 38-bp and 55- to 67-%, respectively (Table 1). The location of the primers and their orientations within the selected target gene in the whole genome of *D. solani* are shown in Figure 1.

### Assay specificity

Among 133 bacterial strains (Supplement Table 1) that were tested to determine the LAMP assay specificity and efficiency, no false-positive or false-negative results were observed, i.e., LAMP results matched 100% with all tested strains of *D. solani* (Figure 2). LAMP amplification shown as a sigmoid shaped curve in the plot produced by the real-time qPCR thermocycler with a distinct melt curve ranging from T_m_ 90.86-91.06, with a T_t_ (threshold time) from 5.78-6.39 mins with all *D. solani* strains tested (Figures not shown). For visual detection protocol, SYBR Green I was adapted, the reaction solution in the tubes with target DNA amplification turned to green color after addition of SYBR Green I and showed high fluorescence under the UV light (Figure 2). A ladder/smear-like pattern was also observed when amplified products were electrophoresed and visualized on a 2% agarose gel, indicating positive amplification (Figures not shown). No exponential curve, no melt curve, no color change, were observed with any non-target bacterial strains (from different geographic locations and hosts) included in the exclusivity panel, healthy control plant DNA and non-template control, which demonstrated no cross-reactivity with the developed assay (Supplement Table 1). Thus, the designed primers were highly specific and did not form any primer-dimers or non-specific products.

**Figure 2:**
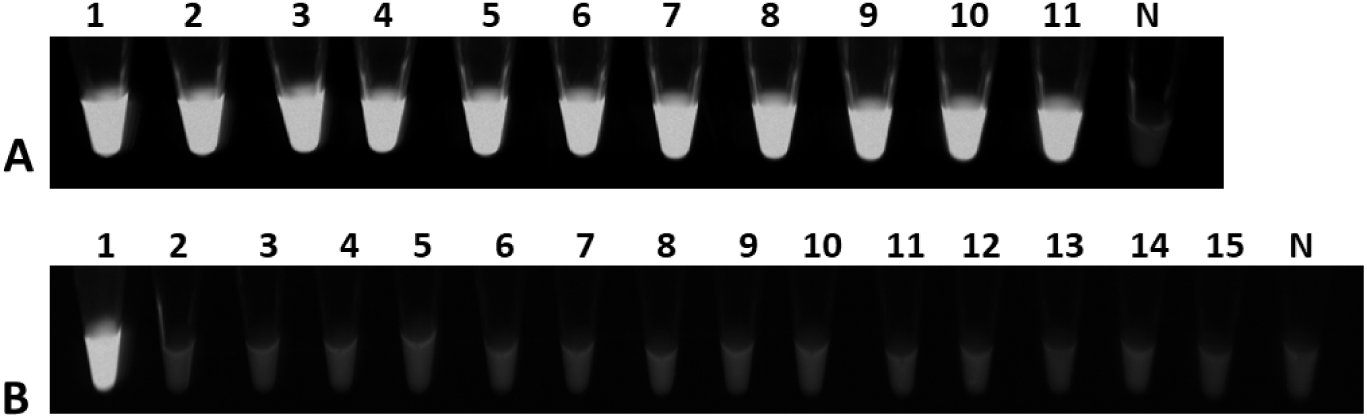
Specificity Validation of Loop-Mediated Isothermal Amplification (LAMP) for *D. solani* Detection. (A, B) Fluorescence detection under UV light. (A) Inclusivity assay with all *D. solani* strains: Tube 1 contains the positive control (genomic DNA of *D. solani* A5581), Tubes 2-11 contain genomic DNA of *D. solani* strains A5582, A6288, A6289, A6291, A6292, and A6294-A6298, and Tube N is the non-template control (NTC; water). (B) Exclusivity assay with genomic DNA from thirteen representative bacterial strains from the exclusivity panel: Tube 1 contains *D. solani* (A5581); Tubes 2–15 contain *D. dianthicola, D. paradisiaca, P. carotovorum, D. zeae, P. brasiliense, D. chrysanthemi, P. carotovorum, Pantoea* sp., *D. dadantii, E. amylovora, Klebsiella* sp., *P. betavasculorum, P. odoriferum*, and healthy *S. tuberosum* (negative control); N is water (NTC).

The specificity of TaqMan-qPCR was first tested *in-silico* by using the BLASTn tool against the NCBI GenBank database, primers showed 100% query coverage and 100 % identity match with the targeted *D. solani,* indicating high specificity of the designed primers. The flap sequences allowed the *D. solani* primer sets to be multiplexed with the previously developed and validated *Dickeya* genus primers. Multiplexing with the previous primers set does not show any cross amplification with non-targets (data not shown). Additionally, when the developed multiplex TaqMan qPCR assay was tested with 123 bacterial strains listed in Supplement Table 1, positive results were only observed in orange and green channels (specific for *D. solani* two target regions) with 11 strains of *D. solani*; and 43 other strains in genus *Dickeya* in yellow channel (Supplement Table 1). The C_T_ values from *D. solani* strains for *LysR* transcriptional regulator gene target ranged from 15.33 ±0.08 to 22.35±0.17 in orange channel (n=11), while the green channel for second target gene (*tetR* family transcriptional regulator gene) exhibited almost similar C_T_ values of 14.89±0.03 to 22.46±0.06 (n=11). The C_T_ value from other *Dickeya* species excluding *D. solani*, ranged from 11.31±0.48 to 33.24±0.31 in yellow channel (n= 46), while *D. solani* strains amplified in range from 14.37±0.01 to 21.56±0.04 (n=11). No amplification or C_T_ values were obtained in yellow, orange or green channels with any members from exclusivity panel that includes phylogenetically closely related species, non-targets, endophytic species sharing close niche (n=65), and healthy control plant DNA and soil samples included in the panel. Moreover, during the specificity validation, no C_T_ value >35 was obtained with any members of inclusivity panels for all primer/prob sets. The positive amplification or C_T_ value was observed in crimson channel with all the reactions containing bacterial strain DNA, healthy control plant DNA and soil indicating no inhibition in the reaction. Addition of 5’ AT-rich flap sequences allowed the multiplexing with the previously designed genus and UIC (universal internal control) primers and demonstrated no negative effect on the specificity of the primers for specific detection of *D. solani*. The standard deviation of three replicates for samples was <1 within the linear range (C_T_ value less than 35), in orange, green or yellow channels except for one sample with standard deviation of 1.31. Consequently, samples above the C_T_ value of 35 were assessed as negative. No amplification signals were detected in healthy potato plants and no-template control (water, NTC). Based on the evaluation of 116 bacterial strains, the multiplex-TaqMan qPCR was 100% specific with no false positive or false negative results, confirming the high specificity. No contradictory results were obtained with either LAMP or multiplex qPCR with any member of the inclusivity and exclusivity panels (Supplement Table 1).

### Assay sensitivity

#### Sensitivity of LAMP Assay

The overnight grown culture of *D. solani* (A5581) confirmed by standard plate counting the serial dilutions of pure cell culture of *D. solani* on the DPA (dextrose peptone agar) media and was found to be 9.5×10^8^ CFU/ml. The detection limit of the developed LAMP assay determined using the ten-fold serially diluted crude cell lysate of pure culture of *D. solani* reached as low as 950 CFU/ml of bacterial cells or ~1 CFU/reaction. A visual detection of the LAMP amplified products after adding SYBR Green I to the LAMP amplified products; fluorescence under UV light is shown in Figure 2.

The T_t_ for detection from 9.5×10^8^ CFU/ml to the lowest limit of detection (950 CFU/ml) was 8.92 min and 5.73 min, respectively. The limit of detection using ten-fold serially diluted genomic DNA of *D. solani* was 100 fg. The sigmoid amplification plot, followed by visual detection by adding SYBR Green I, under UV light and a ladder like amplification from the LAMP amplified products are shown in the Figure 3. Additionally, spiked assays performed by adding 5 μl of dilution buffer prepared from healthy-green-house grown potato plants cell lysate (prepared as described previously using Plant Lysis kit) in LAMP reaction containing ten-fold serially diluted DNA) bacterial crude cell lysate, and 2) genomic DNA to ascertain any inhibitory effect or effect of background host DNA. Similar detection limits of 950 CFU/ml or ~1 CFU/ reaction and 100 fg was observed for the spiked assays performed with the crude cell bacterial lysate and genomic DNA, respectively; indicating no inhibitory effect of the background host crude DNA (Figure 3).

**Figure 3.**
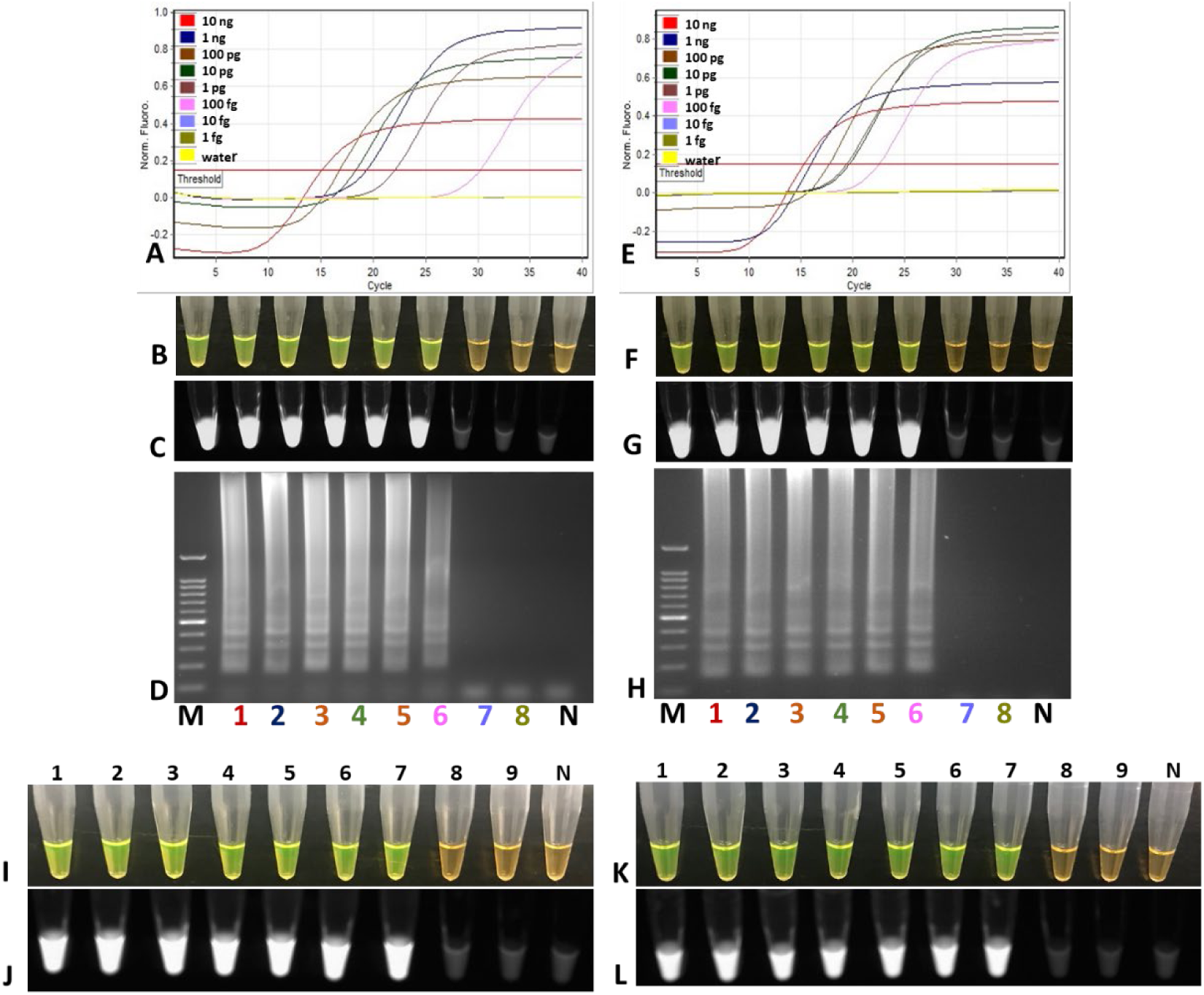
**(A-H)**: Sensitivity assays of loop mediated isothermal amplification (LAMP) for specific detection of *D. solani.* (A-D) LAMP sensitivity assay; Lane M, 100 bp ladder, Lanes 1-8, ten-fold serially diluted purified genomic DNA of *D. solani* from 10 ng to 1 fg.; Lane N, negative template control (NTC, water). (E–H) Spiked LAMP sensitivity assay; lane M, 100 bp ladder; Lanes 1-8 ten-fold serially diluted (1 ng to 1 fg) purified genomic DNA of *D. solani* <u>plus</u> 1 µl of host DNA (10 ng/µl) added to each reaction; Lane N, negative template control (NTC, water). (A & E) Rotor-Gene Q amplification of sensitivity assay and spiked sensitivity assay; (B & F) visual detection of LAMP after adding SYBR Green I; (C & G) tubes under UV; (D & H) amplified LAMP products on 2% agarose gel. **(I-L)**: Visual detection of sensitivity assays of *D. solani* heat-killed cells using LAMP assay. (I, J) Tubes 1-9 ten-fold dilution series consisting of 10^8^, 10^7^, 10^6^, 10^5^, 10^4^, 10^3^, 10^2^, 10, 1 CFU/ml of heat-killed cells of *D. solani*, and a non-template control (tube N). (K, L) Tubes 1-9 ten-fold dilution series consisting of 10^8^, 10^7^, 10^6^, 10^5^, 10^4^, 10^3^, 10^2^, 10, 1 CFU/ml of heat-killed cells of *D. solani* spiked with host lysate, and a no-template control (tube N). Products were detected after adding 3 µl SYBR Green I under visual light, where a positive result changes from orange to green (I, K), or ultraviolet light, where a positive result show fluoresce (J, L).

#### TaqMan-qPCR Sensitivity Assay

The primer and probe sets were compared for their sensitivity in multiplex and singleplex formats, and also in the presence of host DNA background. The *D. solani* primer sets (Dso-wF1/wR1, Dso-wF2/wR2) targeting two different unique loci of the *D. solani* genome, contained 5’ flap sequences, were used to make them compatible with the previously *Dickeya* genus (DICg-wF1/wR1) and internal control primers (UIC-wF/wR). In 4-plex TaqMan-qPCR, *D. solani* specific primer sets, Dso-wF1/wR1 and Dso-wF2/wR2, were able to detect down to 1 pg of genomic DNA with almost similar C_T_ values of 30.38±0 and 29.86±0 (Figure 4) with correlation coefficient (R^2^) of 0.99; and amplification efficiency (E) of 0.99 and 0.94, respectively. The average genome size of *D. solani* is 4.9 Mb. Based on the genome size, copy number was calculate, using the web-based software (http://cels.uri.edu/gsc/resources/cndna.html or http://www.scienceprimer.com/copy-number-calculator-for-realtime-pcr). Based on the formula, 1.89×10^6^ copies were present in 10 ng of genomic DNA. In 4-plex qPCR, detection limit of each primer set specific to detect *D. solani* was down to 1 pg which is equivalent to 180 copies. No difference in the sensitivity was observed when 1 µl of host genomic DNA (DNA extracted from healthy potato plant) was added to each ten-fold serially diluted genomic DNA. The detection limit of 1 pg (180 copies); with almost similar C_T_ values 31.56±0 and 30.13±0 with each correlation coefficient (R^2^) of 0.99; and amplification efficiency (E) of 0.99 and 0.94, respectively. This demonstrates that the developed method is robust with no adverse effect from host background DNA (Figure 4 a and b B1, B2). But, in 3-plex qPCR, performed without addition of UIC in the reaction mix, a ten-fold increase in the detection limit was observed, the sensitivity was down to 100 fg equivalent to 18 copies for Dso-wF1/wR1, Dso-wF2/wR2 each primer sets with almost identical C_T_ 34.78±0 and 33.36±0; with each correlation coefficient (R^2^) of 0.99; and amplification efficiency (E) of 0.99 and 0.96, respectively. The primer sets were also compared by performing single qPCR with only single primer set for each target gene specific for *D. solani*. Each primer set Dso-wF1/wR1 and Dso-wF2/wR2 was able to detect down to 10 fg of *D. solani* genomic DNA with identical C_T_ value 37.26±0 and 37.40±0.39; with each a correlation coefficient (R^2^) of 0.99; and amplification efficiency (E) of 0.99 and 0.98, respectively (Figure 4 a and b, D and E). The reaction efficiencies ranged from 90 to 101%, indicating high linearity. The detection limit of each qPCR primer set specific for *D. solani* was down to 1.8 copies i.e., ~ 2 copies of the genomic DNA in a single-plex qPCR. The sensitivity of the *Dickeya* genus primers in the 4-plex qPCR was 1 pg (180 copies) with C_T_ value 29.86±0 with a correlation coefficient (R^2^) of 0.98; and amplification efficiency (E) of 0.90 (Figure 4 a and b, A3). No change in the sensitivity was observed when spiked assay was performed by adding host genomic DNA with almost identical with C_T_ value 29.33±0 (1 pg), correlation coefficient (R^2^) of 0.99; and amplification efficiency (E) of 0.93 (Figure 4a and b, B3). Similarly, ten-fold increase in sensitivity was observed in 3-plex qPCR without adding UIC in the reaction mix. The primer detection limit was down to 100 fg (18 copies) with C_T_ value 32.24±0, correlation coefficient (R^2^) of 0.99; and amplification efficiency (E) of 0.96 (Figure 4a and b, C3).

**Figure 4:**
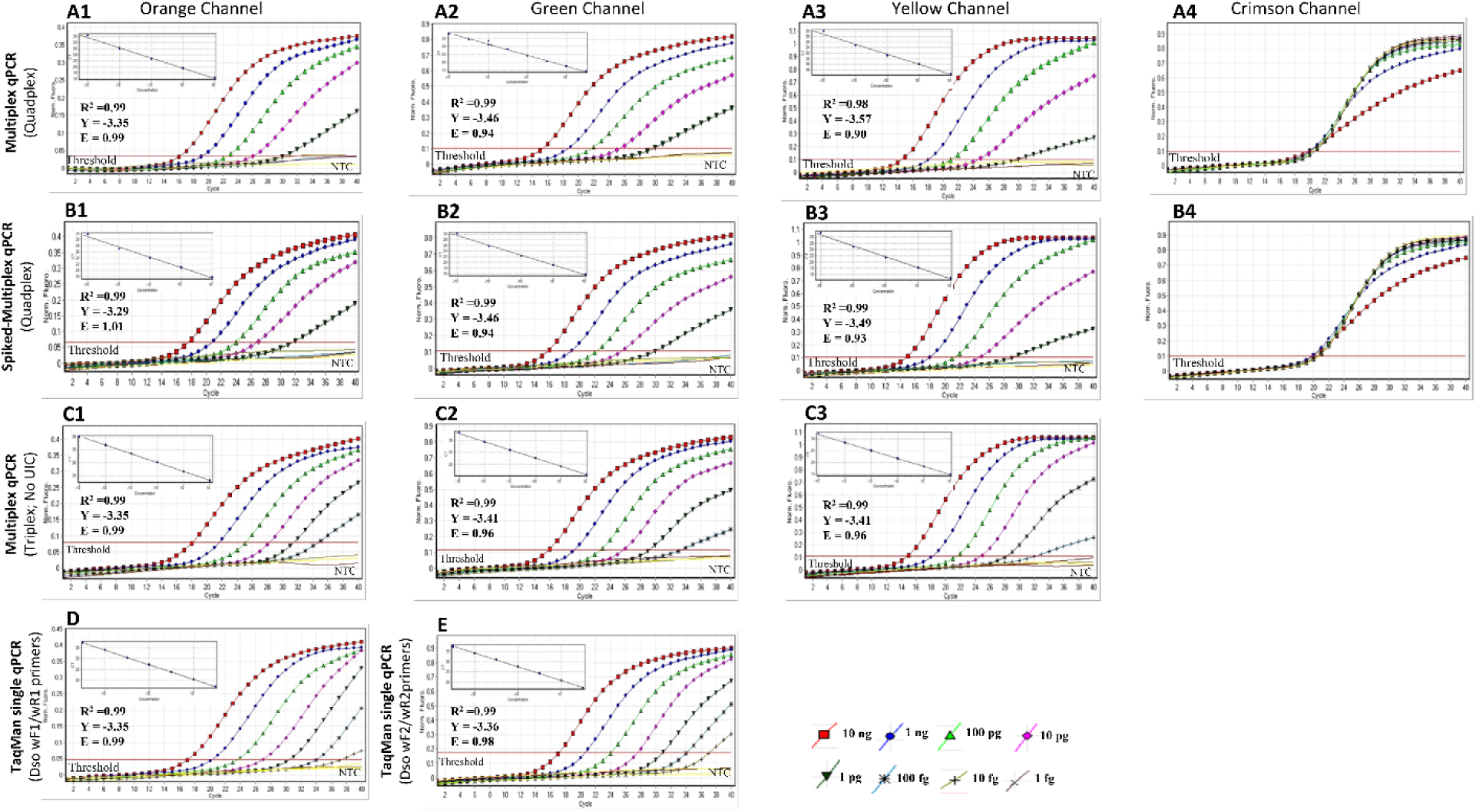
Standard curves and graphs for TaqMan-qPCR generated using ten-fold serial diluted *Dickeya solani* genomic DNA (10 ng to 1 fg). (A, C) Multiplex TaqMan-qPCR, (D, E) Single TaqMan-qPCR with 10-fold serially diluted genomic DNA; and (B) Multiplex TaqMan qPCR 10-fold serially diluted genomic DNA mixed with host plant DNA. The Orange, Green, Yellow and Crimson Channels correspond to the different reported dye Red –BHQ2, 6-FAM (495/520), HEX (535/554), Quasar705-IBQ3 (excitation/emission spectra in nm), respectively. A1/A2/A3/A4 multiplex TaqMan qPCR generated by multiplexing Dso-wF1/wR1/Dso-P1, Dso-wF2/wR2/Dso-P2, DICg-wF1/wR1/DICg-P, UIC-wF/wR/UIC-P primer/probe sets. B1/B2/B3/B4 spiked multiplex TaqMan qPCR generated by multiplexing Dso-wF1/wR1/Dso-P1, Dso-wF2/wR2/Dso-P2, DICg-wF1/wR1/ DICg-P, UIC-wF/wR/ UIC-P primer/probe sets; the spiked assay was done by adding 1 μl of healthy plant-potato DNA extracted from tubers to each 10-fold serially diluted *D. solani* DNA. C1/C2/C3 multiplex TaqMan qPCR were generated by multiplexing Dso-wF1/wR1/Dso-P1, Dso-wF2/wR2/Dso-P2, DICg-wF1/wR1/DICg-P primer/probe sets. D single TaqMan qPCR with Dso-wF1/wR1/Dso-P1 primer set in the reaction mix. E single TaqMan qPCR with Dso-wF2/wR2/Dso-P2 probe and primer set. X axis represents the number of cycles, and Y axis-normalized fluorescence. The C_T_ values are average of three replicates ±SD. Slopes (Y = threshold cycles (Ct) of target DNA detected), R^2^ (correlation coefficient), and E (amplification efficiency).

### Assay specificity with artificially and naturally infected samples

Naturally and artificial inoculated “disease free” potato tubers were used to assess the capability of the developed LAMP and TaqMan qPCR assay. No cross-amplification was observed with genomic DNA from all 10 naturally infected potato plants which included PL68 (PS32F), PL109 (PS33), PL107 (PS60) (*P. brasiliense*); PL124 (PS38), PL75 (PS63) (*P. parmentieri*); PL125 (PS1), PL127 (PS10), PL122 (PS66) (*D. dianthicola*); PL66 (PS2) (*P. aroidearum)*; PL73 (*P. carotovorum*) and healthy potato plant samples (Figure 5A). Amplification sigmoidal plot with a melt curve T_m_ 90.46 was only observed with the positive control (Figure 5B).

**Figure 5.**
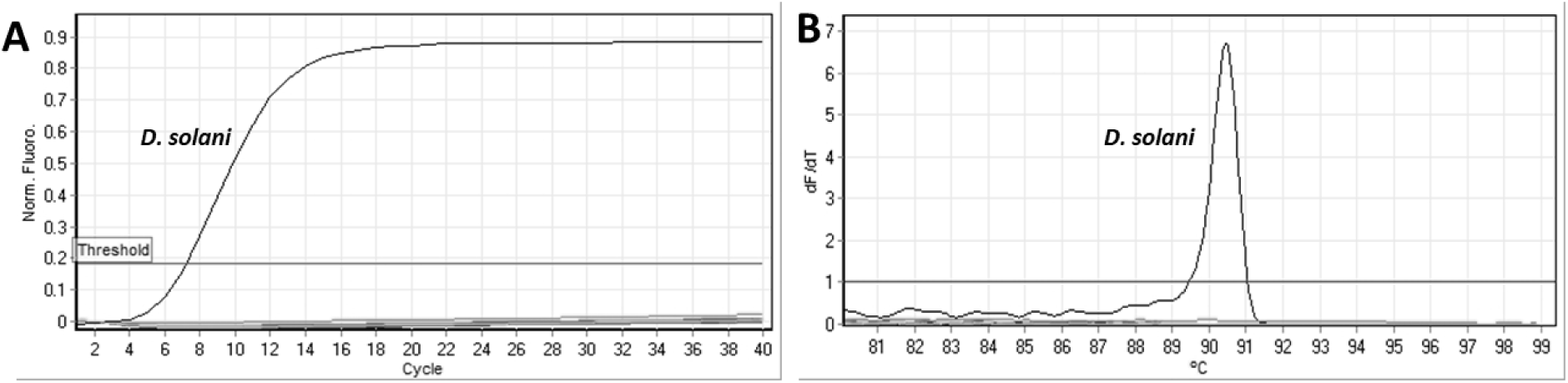
Validation of *Dickeya solani* specific loop mediated isothermal amplification (LAMP) assay with plant samples naturally infected with other soft rot bacteria. *Pectobacterium. brasiliense* PL68 (PS32F), PL109 (PS33) and PL107 (PS60); *P. parmentieri* PL124 (PS38), and PL75 (PS63); *D. dianthicola* PL125 (PS1), PL127 (PS10), and PL122 (PS66); *P. aroidearum* PL66 (PS2); *P. carotovorum* PL73 and healthy potato plant sample. (A) Real time amplification plot, (B) Melting curve. A positive control includes DNA from *D. solani* (A6295; positive amplification shown in both figures).

Additionally, artificially inoculated potato tubers tested with the LAMP showed positive amplification plot within 3.20-3.86 min with only *D. solani* strain A5582, A6289, A6291, A6292, A6295-A6298 infected crude plant cell DNA and no amplification was observed with crude plant cell DNA from *D. dianthicola* (PL25) and *D. paradisiaca* (A6064) and healthy potato. Positive and negative template control (NTC, water) were also included in the run (figure not shown). A visual detection by adding SYBR Green I stain with a change of color from orange to green for positive amplification and no change in color indicating no amplification; the fluorescence under UV light have been shown in Supplement Figure 1. The developed LAMP assay was able to detect 0% false-positive, with 100% efficiency and accuracy, with all samples tested with no cross-amplification with the non-targets.

Table 3 summarizes the results of the developed multiplex TaqMan-qPCR assay, which includes 8 naturally infected potato plants and 9 artificially inoculated tubers. Detection from artificially inoculated tubers showed only amplification with all *Dickey*a species with genus specific primers (yellow channel) with C_T_ values in range from 17.47±0.05 to 25.76±0.99. Both *D. solani* specific primers sets Dso-wF1/wR1, Dso-wF2/wR2 showed positive amplification with *D. solani* infected samples in orange and green channel, with C_T_ values in range from 21.43±0.05 to 29.49±1.06 and 18.23±0.04 to 26.44±0.97, respectively. No amplification was observed with *D. dianthicola* infected and, healthy plant controls or negative template control. Positive amplification was observed in crimson channel with internal control, indicating no inhibition in the reaction. Similarly, amplification was observed with naturally infected samples, with positive amplification in yellow channel for “Genus” with only *Dickeya* species with C_T_ values ranged from 17.93 ±0.03 to 29.80±0.04. No amplification was observed with *D. dianthicola-* and *Pectobacterium* species-infected samples in channels specific for *D. solani*, confirming the specificity of the assay. No amplification was observed with the healthy plant DNA. The developed multiplex-TaqMan qPCR accurately detected, with 100% efficiency, all infected plant samples tested. No false positives or false negatives were detected (Table 3). No conflicting results were observed with all samples tested with both developed methods.

**Table 3.** Validation of 4-plex TaqMan qPCR assay with naturally and artificially infected plant materials.

| Strain | Sample Type | Strain/Sample Number | TaqMan qPCR |  |  |  |
| --- | --- | --- | --- | --- | --- | --- |
|  |  |  | Mean Ct (SD) Rotor Gene |  |  |  |
|  |  |  | Yellow Channel<br>(Genus) | Orange Channel<br>(Dso-P1) | Green Channel<br>(Dso-P2) | Crimson Channel<br>(UIC) |
| Naturally infected plant samples |  |  |  |  |  |  |
| *Dickeya solani* | Bacterial genomic DNA | LMG27550 | 17.93 (0.03) | 20.29 (0.05) | 19.65 (0.05) | 28.91 (0.47) |
| *Pectobacterium brasiliense* | Naturally infected plant | PS32 | - | - | - | 28.03 (0.14) |
| *P. brasiliense* | Naturally infected plant | PS33 | - | - | - | 28.25 (0.34) |
| *P. parmentieri* | Naturally infected plant | PS38 | - | - | - | 28.58 (0.38) |
| *P. brasiliense* | Naturally infected plant | PS60 | - | - | - | 28.41 (0.18) |
| *D. dianthicola* | Naturally infected plant | PS1 | 27.95 (0.12) | - | - | 28.45 (0.17) |
| *D. dianthicola* | Naturally infected plant | PS10 | 20.29 (0.04) | - | - | 27.75 (0.12) |
| *D. dianthicola* | Naturally infected plant | PS66 | 25.52 (0.11) | - | - | 27 (0.19) |
| *P. aroidearum* | Naturally infected plant | PS2 | 29.80 (0.04) | - | - | 28.15 (0.35) |
|  | Healthy potato | NC | - | - | - | 28.53 (0.44) |
|  | water | NTC | - | - | - | 27.67 (0.69) |
| Artificially inoculated tubers |  |  |  |  |  |  |
| *D. solani* | Bacterial genomic DNA | LMG27550 | 17.47 (0.05) | 21.43 (0.05) | 18.23 (0.04) | 29.45 (0.40) |
| *D. solani* | Artificially inoculated | LMG25865 | 25.76 (0.99) | 29.49 (1.06) | 26.44 (0.97) | 27.56 (0.35) |
| *D. solani* | Artificially inoculated | LMG25990 | 24.72 (0.0) | 28.50 (0.0) | 25.46 (0.0) | 26.89 (0.0) |
| *D. solani* | Artificially inoculated | LMG25993 | 24.08 (0.43) | 27.86 (0.31) | 24.95 (0.42) | 27.15 (0.27) |
| *D. solani* | Artificially inoculated | LMG27552 | 23.09 (1.52) | 26.88 (1.48) | 23.89 (1.55) | 27.77 (0.22) |
| *D. solani* | Artificially inoculated | LMG27553 | 24.84 (0.0) | 28.61 (0.0) | 25.64 (0.0) | 27.82 (0.0) |
| *D. solani* | Artificially inoculated | LMG27554 | 23.93 (0.06) | 27.74 (0.04) | 24.69 (0.09) | 27.34 (0.05) |
| *D. solani* | Artificially inoculated | A5582 | 23.97 (0.03) | 27.82 (0.08) | 24.71 (0.02) | 27.26 (0.14) |
| *D. solani* | Artificially inoculated | LMG27550 | 23.54 (0.0) | 27.27 (0.0) | 24.39 (0.0) | 26.76 (0.0) |
| *D. dianthicola* | Artificially inoculated | PL25 | 24.27 (0.08) | - | - | 28.06 (0.54) |
|  | Healthy potato | NC | - | - | - | 28.46 (0.25) |
|  | water | NTC | - | - | - | 28.26 (0.21) |

\*Ct threshold time for detection; SD – standard deviation

### Multi-laboratory validation of LAMP and multiplex-TaqMan qPCR Assays

No false positive and/or false negative results were observed with eight bacterial DNA that were blind-coded across the two-testing laboratory in the multi-lab validation. Same samples were used to test with the optimized LAMP and multiplex-TaqMan qPCR. Table 4 summarizes the results of the LAMP and multiplex-TaqMan qPCR. The developed LAMP assay was able to detect with 100% accuracy and efficiency with amplification only with the *D. solani* genomic DNA and no amplification with *D. dianthicola*, *D. zeae* and healthy control plant DNA included in the panel. No false positive or false negatives were detected during the run in two labs tested with two different machines.

**Table 4.**
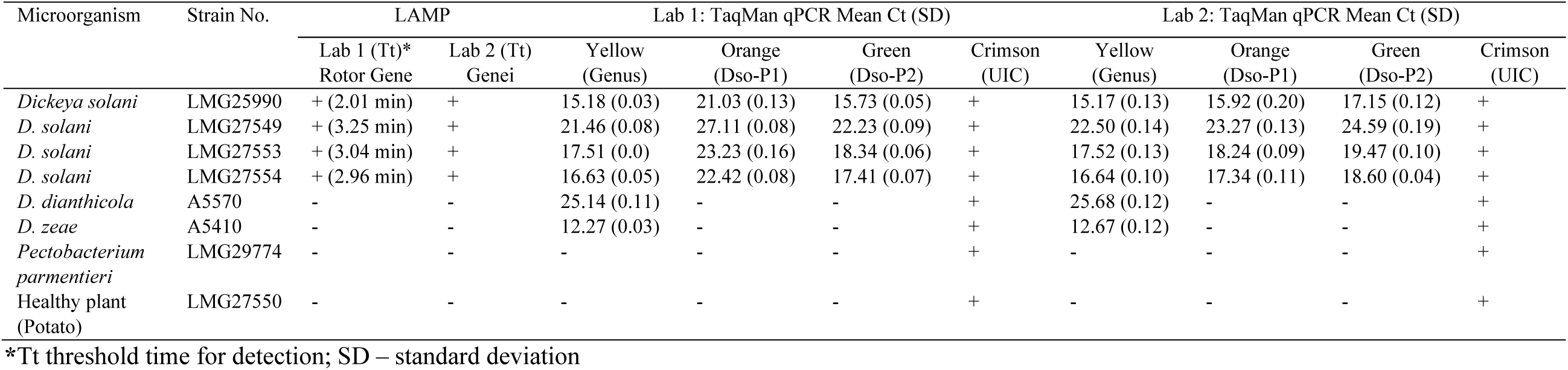
Multi-laboratory validation of loop-mediated isothermal amplification (LAMP) and TaqMan qPCR for specific detection of *Dickeya solani*.

Again, same samples were tested with the multiplex TaqMan-qPCR, the C_T_ values were obtained in the different channels in two separate labs using different machines (Table 4). The C_t_ value ranging from 12.27±0.03 to 25.14±0.11 (lab 1) and 12.67±0.12 to 25.68±0.12 (lab 2) was observed with all *D. solani-* and *D. zeae-*infected samples’ DNA using genus specific primer/probe set (DICg-wF1/wR1/DICg-P). Amplification was observed with both primer sets designed for specific detection of *D. solani,* and no cross-reaction was observed. Mean Ct values for *D. solani* Dso-wF1/wR1/Dso-P1 (orange channel) primer/probe set ranged from 21.03±0.13 to 27.11±0.08 (lab 1) and 15.92±0.20 to 23.27±0.13 (lab 2). Mean Ct values for *D. solani* Dso-wF2/wR2/Dso-P2 (green channel) primer/probe set ranged from 15.73±0.05 to 22.23±0.09 (lab 1) and 17.15±0.12 to 24.59±0.19 (lab 2). Furthermore, positive amplification was observed with universal internal control with all samples tested indicating no inhibitory effect in the reaction. The samples tested in both labs produced identical results with 100% accuracy and efficiency indicating the developed assay is robust and reliable for testing the infected samples.

## Discussion

*Dickeya solani* stands out as one of the most aggressive causative agents of blackleg and soft rot, thrives across various temperatures, outperforming its genus counterparts in spreading disease (Heuer et al., 2010; Czajkowski et al., 2013; Potrykus et al., 2014). Prevalent in Europe and Israel, it spreads through the trade of latently infected seed potatoes. Although not yet documented in the United States, *D. solani* is a quarantine pathogen due to the significant risk it poses to the potato industry. The extensive germplasm trade makes its introduction through imported seed potatoes almost inevitable, highlighting the need for vigilant monitoring and control measures.

This research presents sensitive and accurate LAMP and TaqMan qPCR methods for detecting *D. solani*. These methods, targeting unique genomic regions of *D. solani*, have been thoroughly validated, offering flexibility to suit various lab needs. The TaqMan-qPCR’s primers and probes are versatile, usable in both single and multiplex formats, catering to user preferences. The multi-gene-based approach is reported to enhance reliability, accuracy, and specificity while minimizing the risk of false positives (Arif et al., 2013). The dual target-based probes in real-time qPCR provides additional advantage and high specificity and reliability of the assay. *In vitro* validation, using strains that represent various geographical locations, hosts, and closely related taxa, is crucial for developing a reliable diagnostic assay (Dobhal et al., 2018; Arizala et al., 2022). Both the LAMP and multiplex-TaqMan-qPCR assays underwent extensive validation using a diverse range of bacterial strains collected from various geographical locations and both artificially and naturally infected plant samples, ensuring high accuracy and reliability. The assays were also tested against saprophytic and endophytic bacterial strains from similar niche/environments, as well as infested soil. Both the LAMP and multiplex-TaqMan-qPCR assays exhibited high specificity, delivering 100% accurate results with the inclusivity and exclusivity panels. No false positives or false negatives were detected during validation, affirming the reliability and robustness of the developed assays.

Diagnostic assays targeting unique genomic regions offer robustness and high accuracy (Boluk et al., 2020; Dobhal et al., 2020; Arizala et al., 2022). A comparative whole-genome approach was utilized to identify conserved and unique genomic regions (*lysR* and *tetR* gene regions). These target genes were used in designing specific, thermodynamically competent primers and probes for *D. solani*. For the LAMP assay, primers were designed based on the *t*etR family transcriptional regulator gene. *In-silico* analysis of these primers using NCBI BLASTn confirmed their high specificity, showing 100% identity match and 100% query coverage for *D. solani*. Moreover, *in-vitro* testing with DNA isolated from various type strains and reference strains from international and national culture collections demonstrated 100% specificity of the assays, with no occurrences of false positives or negatives.

Currently, no LAMP assay has been developed specifically for detecting *D. solani*. A LAMP assay for detecting the *Dickeya* genus, reported by Yasuhara-Bell et al. (2017), offers a sensitivity of 5 pg/reaction and a detection time of 40 minutes per reaction. However, the LAMP assay for *D. solani* developed in this research is more sensitive and rapid, detecting as little as 100 fg/reaction, with results obtainable in under 9 minutes (the quickest detection time in validation was 8.92 minutes). Additionally, when using crude cell lysate, the LAMP assay can detect as few as 950 CFU/ml in a sample, including plant samples, which is equivalent to 1 CFU/reaction. Recently, an isothermal method for *D. solani* was reported by Ivanov et al. (2020), combining a lateral flow assay with recombinase amplification, demonstrating a sensitivity of 700 CFU/ml of the sample with results detectable in 30 minutes. The rapid LAMP assay presented in this research has been validated with extensive inclusivity and exclusivity panels, including strains from closely related niches.

The LAMP assay introduced in this study stands out for its speed (detection ranging from 5.73 min for the highest concentration to 8.92 min for the lowest) and heightened sensitivity compared to existing methods. This significantly shortens the detection time for the pathogen without sacrificing efficiency or sensitivity. Notably, the LAMP assay demonstrates superior sensitivity (100 fg, equivalent to 18 copies of genomic DNA) when compared to the TaqMan-qPCR assay, which has a detection limit of 1 pg (180 copies) for the 4-plex and was similar to the 3-plex TaqMan-qPCR without the universal internal control (100 fg ~18 copies). The LAMP’s ability to detect crude cell lysate was great, with a sensitivity of 950 CFU/ml, equivalent to 1 CFU/reaction, with a detection time of 8.41 min. LAMP assays are recognized for their robustness, owing to their resilience against inhibitory substances in sample matrices, attributed to the inhibitor-resistant nature of *Bst* polymerase (Poon et al., 2006; Li et al., 2011, Arif et al, 2021). In our LAMP assay the method for preparing crude DNA extract was minimal yet sufficient for amplification, without impacting the detection limit.

Existing literature reports two TaqMan qPCR assays for general *Dickeya* species detection, using *D. solani* bacterial DNA in plant samples, with a sensitivity of 100 fg/reaction (Zijlstra et al., 2020). In our newly developed primer/probe pairs DsowF1/wR1/Dso-P1 and DsowF2/wR2/Dso-P2 detecting as low as 10 fg (~1.8 copies). The 3-plex TaqMan-qPCR, excluding the UIC, and the 4-plex, which includes *Dickeya* genus and UIC primers, have sensitivity of 100 fg and 1 pg/reaction, respectively. Adding the UIC to DsowF1/wR1/Dso-P1 and DsowF2/wR2/Dso-P2 primers and the previously reported *Dickeya* genus primer impacted sensitivity, resulting in a tenfold decrease. This contrasts with Ramachandran et al. (2021), who observed no effect of UIC on sensitivity, suggesting potential incompatibility of UIC primers/probes with our *D. solani*-specific primers. Testing the primers in a 3-plex format maintained a sensitivity of 100 fg for each target gene, including the *Dickeya* genus. However, Dobhal et al. (2020) reported sensitivity of 10 fg per reaction (Ct = 33.75±0.24) with genus *Dickeya*-specific primers. This may be due to compatibility issues or an inhibitory effect among the primer pairs and probes in the 3-plex compared to the 2-plex. The 5’ flap sequences were incorporated into each primer set to improve thermodynamics and to synchronize the reaction, facilitating the combination of primer/probe sets for new multiplex reactions (Arif and Ochoa-Corona, 2013; Larrea et al., 2019).

Detecting the pathogen in infected samples is crucial to demonstrate the developed assay’s practicality (Arizala et al., 2022). Both LAMP and qPCR assays proved their effectiveness in identifying the target pathogen in infected plant samples. The LAMP method showcased 100% specificity, detecting the pathogen accurately in 10 naturally and 10 artificially infected plant tissues, with no false positives or negatives. Similarly, the TaqMan-qPCR assay achieved 100% specificity in identifying the pathogen in 8 naturally infected potato plants and 9 artificially inoculated tubers. In a multi-laboratory validation, both assays consistently delivered 100% concordant results across different labs, underscoring their robustness, specificity, reproducibility, and accuracy.

In summary, we present sensitive and rapid diagnostic assays, including LAMP and multiplex TaqMan-qPCR, developed through a comparative genomics approach to detect *D. solani*, a quarantined, economically impactful bacterial plant pathogen. The LAMP method, resistant to common plant sample inhibitors, is ideal for rapid, on-site detection in fields or at entry ports, crucial for monitoring latent infections in potato seed tubers. Its simplicity makes it suitable for use in basic laboratories. Additionally, the multi-gene-based TaqMan-qPCR, enhanced by including *Dickeya* genus and UIC primers, offers increased confidence and reliability. These tools significantly aid phytosanitary agencies in their swift and effective response, helping to prevent the spread of the pathogen.

## Competing interests

The authors declare that they have no competing interests.

## Funding

This work was supported by the U.S. Department of Agriculture, Animal and Plant Health Inspection Service Farm Bill (APP-7058), USDA National Institute of Food and Agriculture, and Hatch project 9038H managed by the College of Tropical Agriculture and Human Resources. The strains used in this study were revived and maintained with NSF grant support (NSF-CSBR grant no. DBI-1561663). We would also like to acknowledge the culture collections BCCM/LMG (Belgian Co-ordinated collections of Micro-organisms, Belgium), CIRM-CFBP (CIRM-Plant Associated Bacteria, France) and ICMP (International Collection of Microorganisms from Plants, New Zealand) for the strains used for the validation. The mentioned trade names or commercial products in this publication does not imply recommendation or endorsement by the University of Hawaii. The findings and conclusions in this publication are those of the authors and should not be construed to represent any official USDA or U.S. Government determination or policy.

## Supplement Materials

**Supplement Figure 1:**
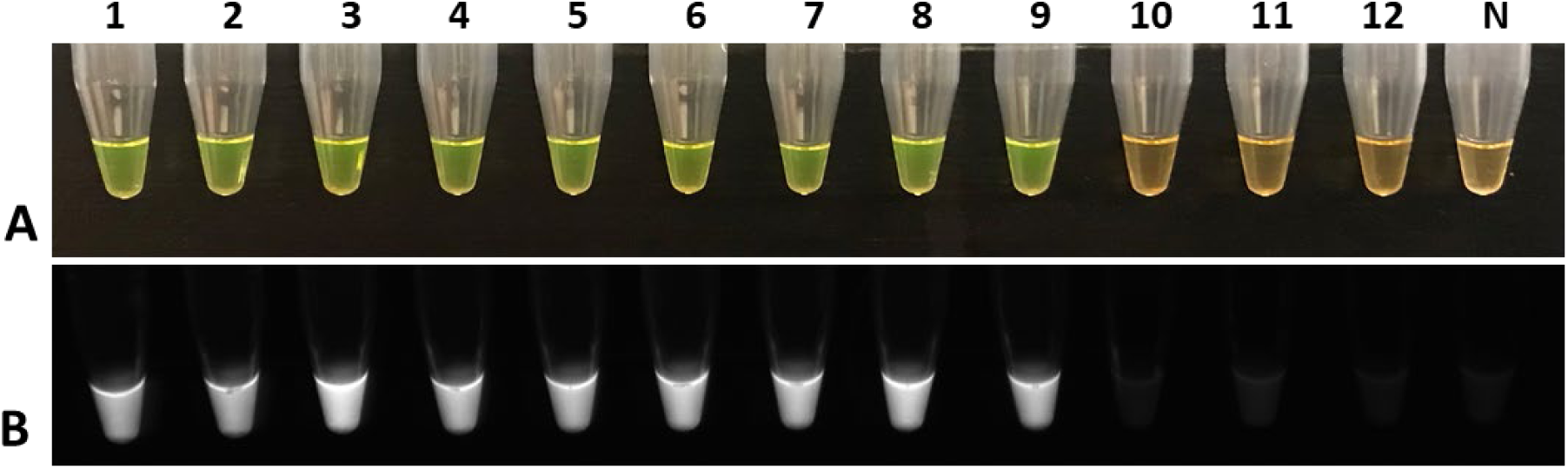
Validation of loop mediated isothermal amplification (LAMP) assay for specific detection of *D. solani* from artificially inoculated “disease-free potato” tubers infected plant samples. (A) Visual observation of LAMP results after adding SYBR Green dye in the amplified LAMP products; (B) LAMP results under UV light. Tube 1 is a positive control (genomic DNA of *D. solani* A6295), tubes 2-9 are tubers inoculated with *D. solani* strain A5582, 6289, A6291, A6292, A6295-A6298, tuelbe 10-11 tuber inoculated with *D. dianthicola* (PL25) and *D. paradisiaca* (A6064), respectively; tube 12 is healthy potato tuber, and tube N is a non-template control (NTC; water).

**Supplement Table 1:**
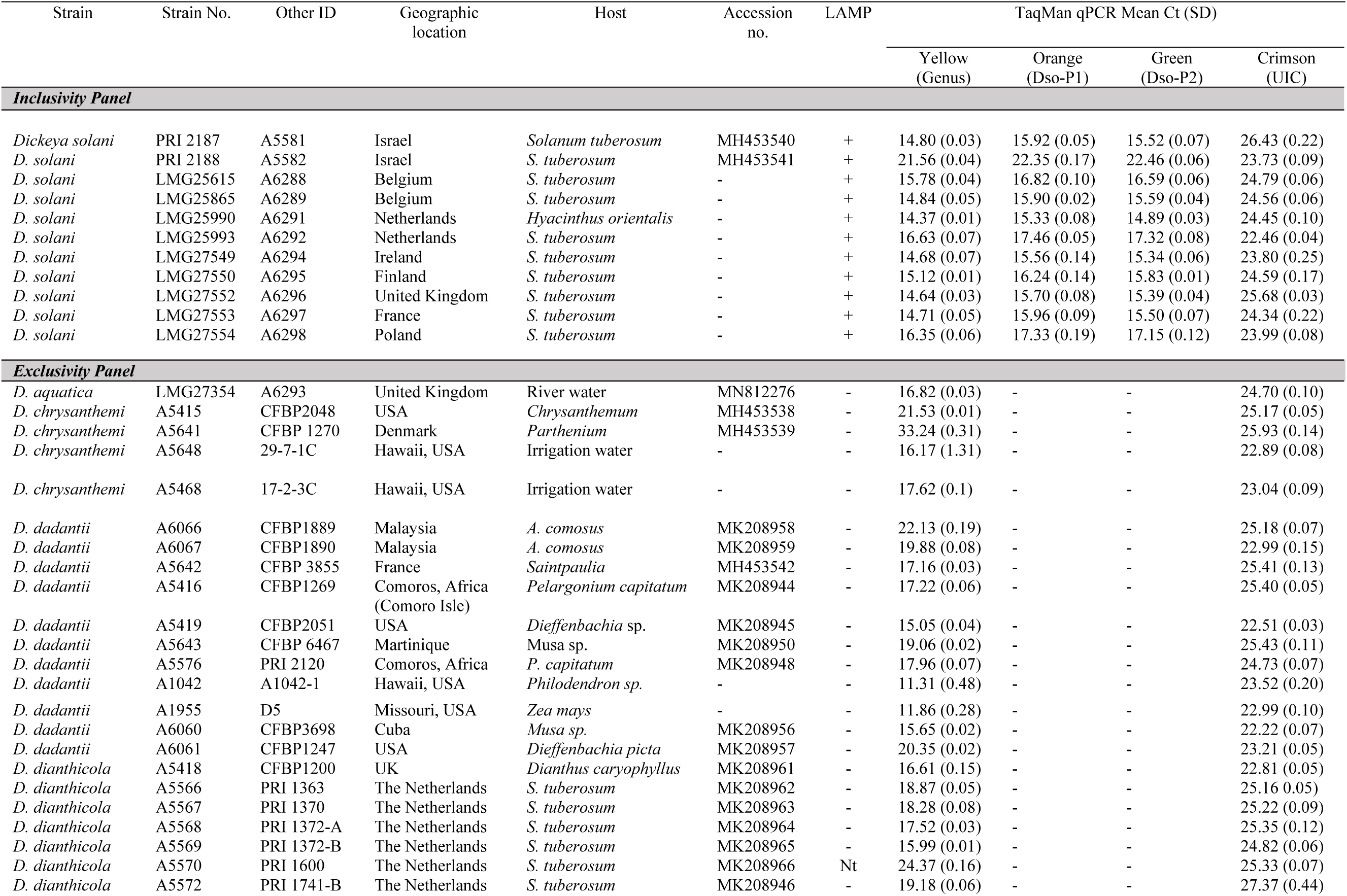
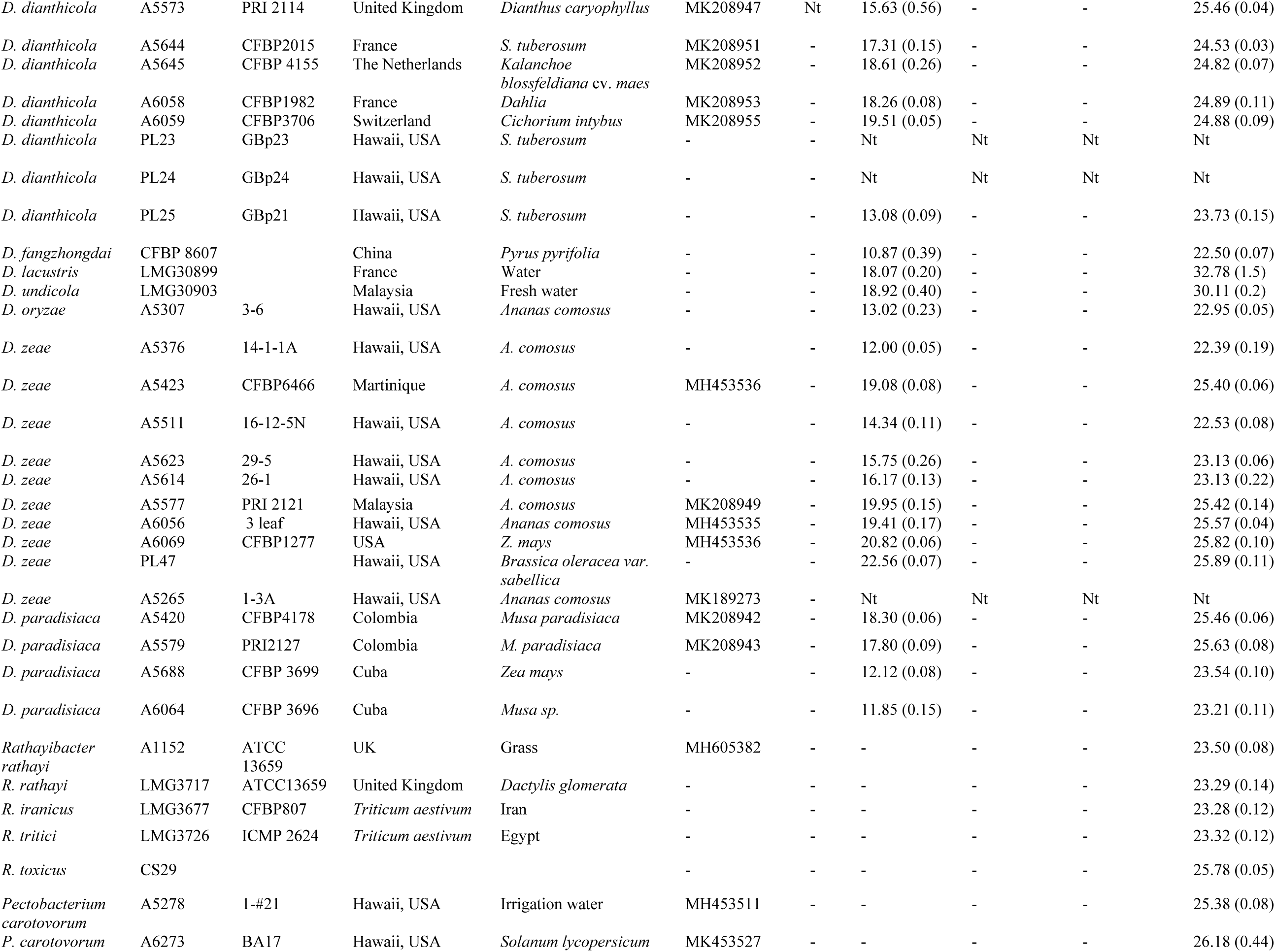
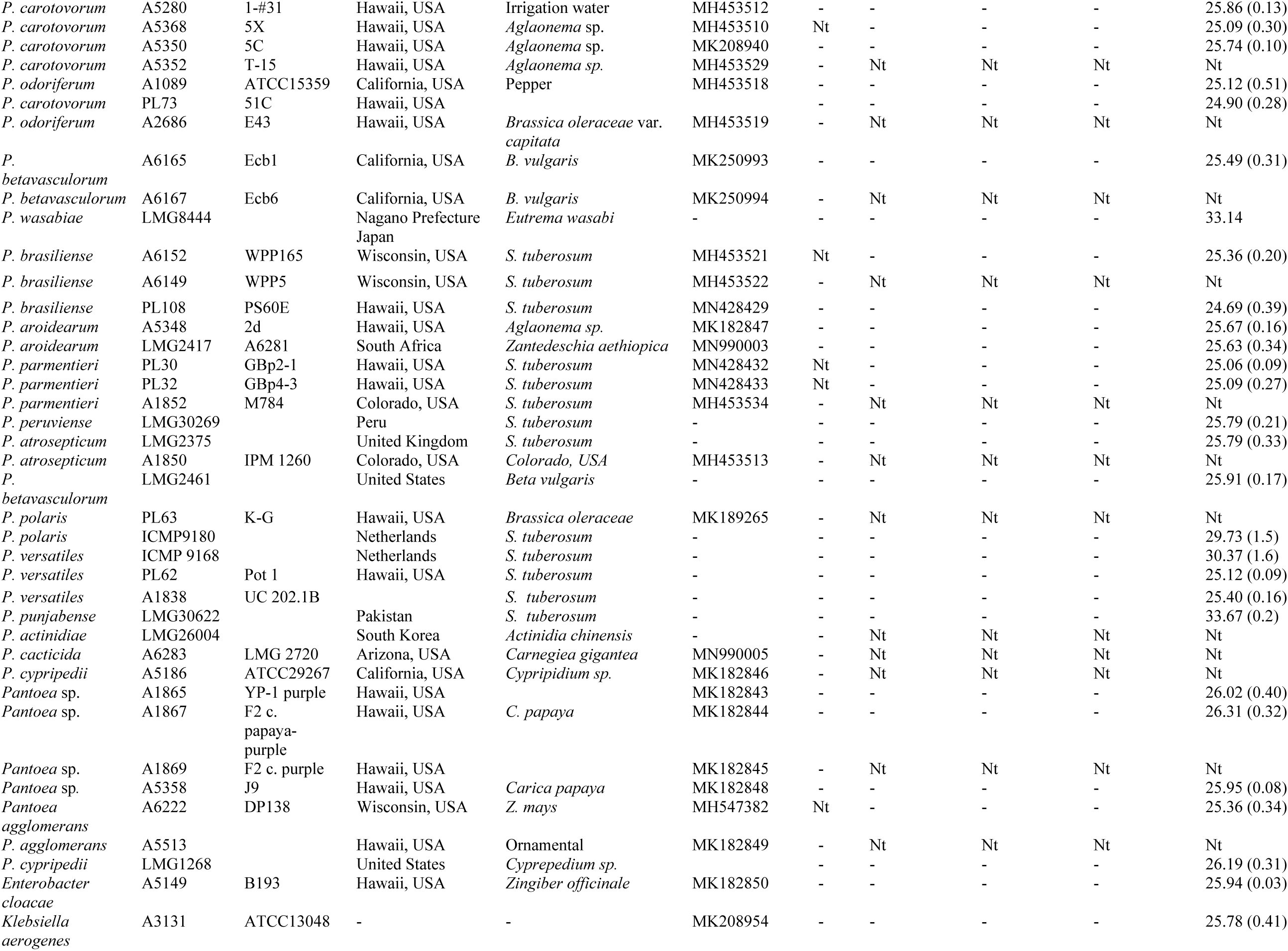
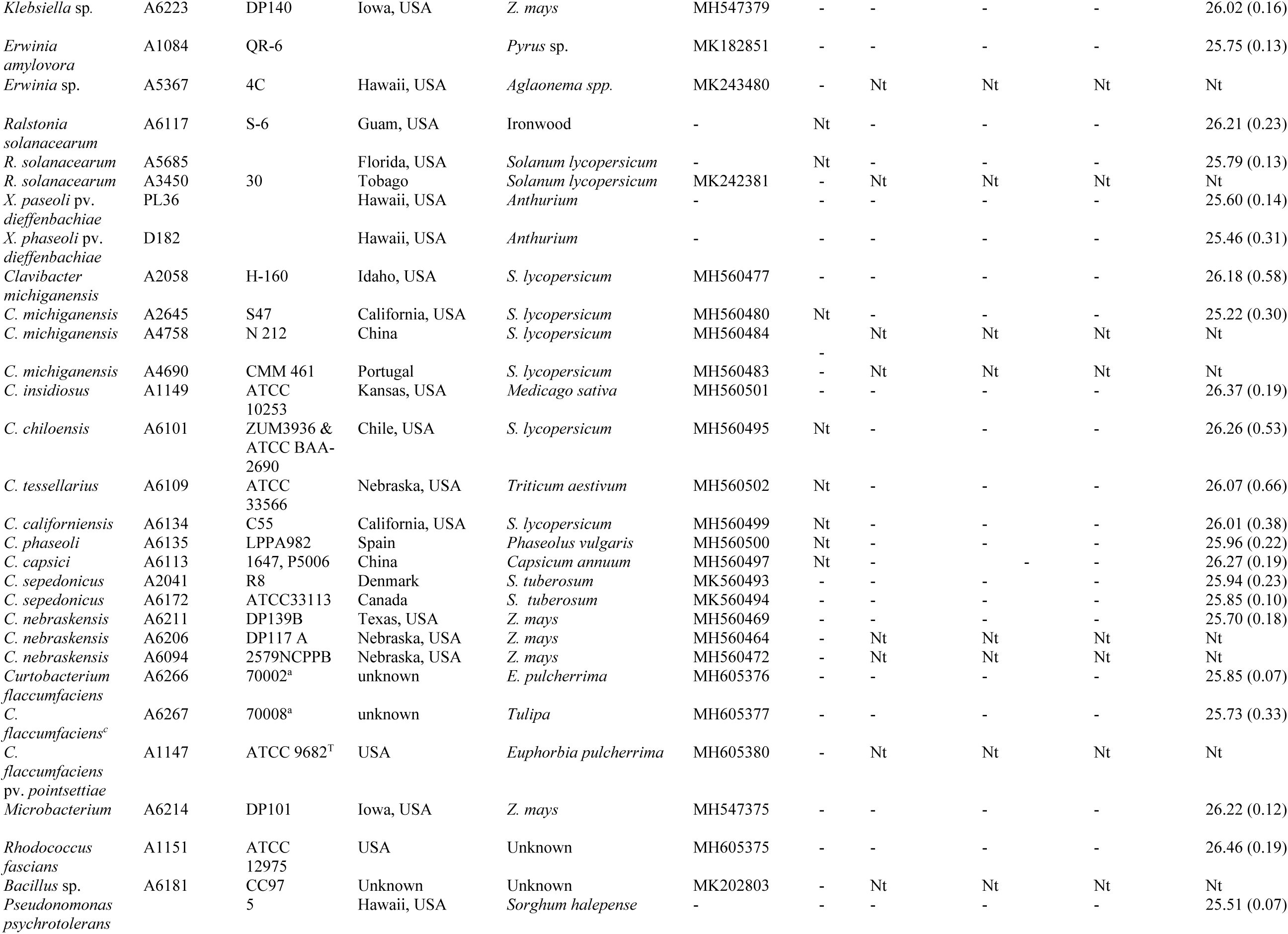
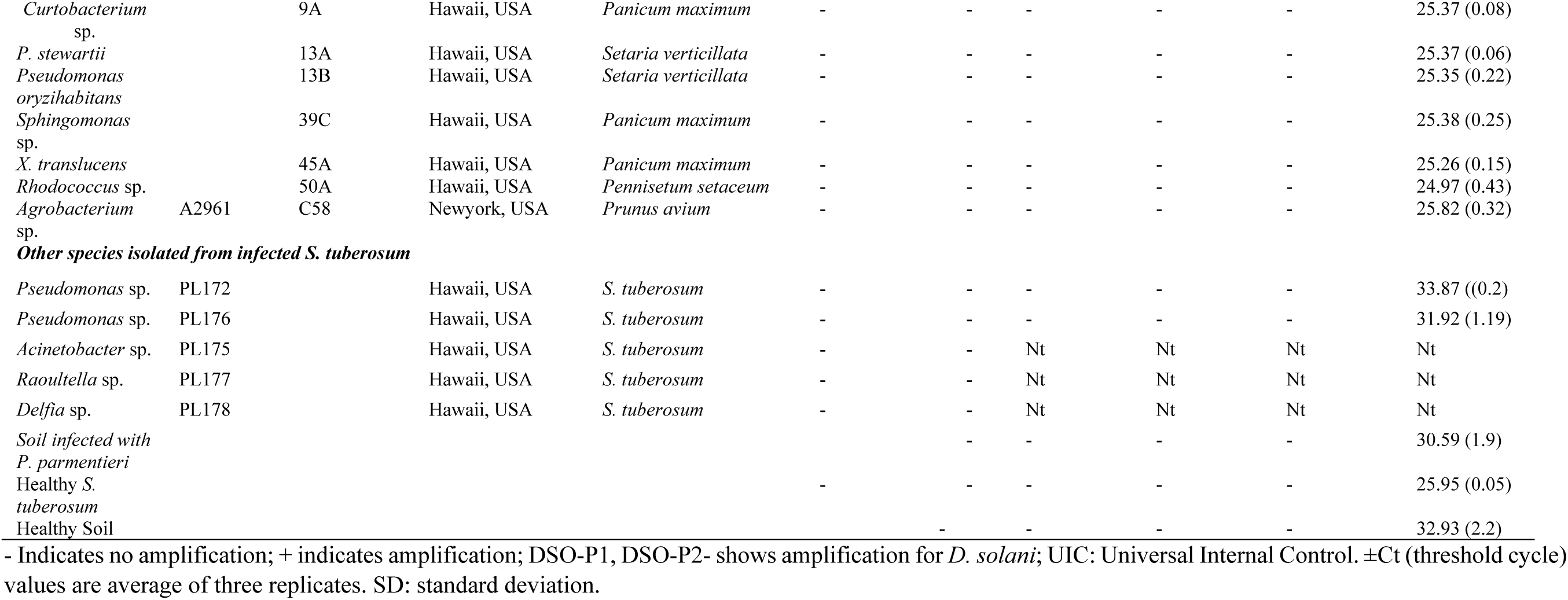
Bacterial strains used for validation of loop mediated isothermal amplification (LAMP) and multi-gene based multiplex TaqMan real-time qPCR for specific detection of *Dickeya solani*.

## Notes

### Competing Interest Statement

The authors have declared no competing interest.

